# Beyond air-seeding: Dynamic, multiphase interactions reveal a two-step mechanism of embolism propagation in angiosperm xylem

**DOI:** 10.64898/2026.08.05.743004

**Authors:** S. Jansen, L. Kaack, M.A. Ahmed, B. Beikircher, P. Bittencourt, R. Chen, J. Flexas, S.M. Gleason, A. Guha, P. Heuret, T. Hölttä, S. Ingram, X. Jiang, P. Jotan, R. Jupa, M. Kanduč, O. Korhonen, M.M. Kotowska, S. Kreinert, A. Laurén, F. Lens, S. Levionnois, R. Link, A. Lintunen, S. Mayr, S.A.M. McAdam, K. Mehltreter, J. Michaud, T.M. Miranda, K. Mocko, P.K. Mondal, H. Morris, A. Nardini, J. Ott, S.S. Paligi, G.S. Pires, L. Plavcová, R.V. Ribeiro, I. Rimer, S. Rosner, L. Rowland, L. Sack, Y. Salmon, A. Scheire, J. Schepers, E. Schneck, B. Schuldt, L.M. Silva, M. Svátek, M. Tomasella, C. Trabi, T. Vesala, R. Welti, Y. Zhang, R. Zweifel, H.J. Schenk

## Abstract

**Background:** The mechanism underlying drought-induced embolism in angiosperm xylem has been attributed to air-seeding. This concept describes the bulk flow of gas from embolised to neighbouring conduits through the penetration of gas-liquid menisci across pores in interconduit pit membranes. While there is compelling evidence for the spatial propagation of embolism, air-seeding rests on various simplifying assumptions. Among others, air-seeding presumes that xylem sap has physical properties comparable to pure water, that pit membranes can be approximated as structures with simple pores, and that embolism occurs whenever a gas-liquid interface crosses a pit membrane.

**Scope:** Recent experimental and theoretical work demonstrates that the biophysical conditions and processes governing gas-liquid interactions at interconduit pit membranes are fundamentally more dynamic and complex than assumed by air-seeding. These phenomena include: (1) gas movement through constriction pore networks, (2) insoluble, polar lipids at conduit surfaces and interfaces, (3) dynamic surface tension of xylem sap that depends on the local packing density of interfacial lipids, (4) bubble snap-off dynamics within pit membranes, (5) surfactant-stabilized nanobubbles in sap that is oversaturated with dissolved gas, and (6) electrostatic interactions between charged interfaces. Importantly, embolism propagation involves bubble generation and embolism formation as distinct, temporarily and spatially separated processes. Embolism formation occurs when nanobubbles become unstable, whereas nanobubbles below critical stability thresholds can remain stable in sap-filled conduits.

**Conclusions:** Together, these findings reconfirm that pit membranes function as safety valves enabling water transport according to the cohesion-tension theory, and provide mechanistic insights into embolism propagation. They address the question why plants do not suffer constant embolism formation despite negative xylem pressures. We conclude that a revised framework explicitly accounting for the 3D structure of pit membranes, and multiphase, dynamic processes operating within them are required to explain the biophysics underlying water transport and embolism resistance in angiosperm xylem.

## Background – a brief historical perspective on air-seeding

A longstanding question in plant biology is how plants transport water under negative pressure without frequent failure by embolism formation (Brown, 2013). Although the cohesion-tension theory (Askenasy, 1895; Dixon and Joly, 1895) is well supported, it does not explain why xylem sap remains stable under tension (Jansen and Schenk, 2015). Air-seeding, a concept coined by Zimmermann (1983), provided one of the first mechanistic explanations for embolism propagation by proposing that gas enters sap-filled conduits from adjacent embolised conduits through interconduit pit membranes. Given the crucial role of long-distance water transport in plants to life on this planet, especially with respect to photosynthesis and plant-climate interactions (Schlesinger and Jasechko, 2014), many may be surprised to hear that our functional understanding of water transport and the mechanisms underlying drought-induced embolism remain a matter of continuing debate and research (Rockwell *et al*., 2014; Lens *et al*., 2022).

The possibility of air-seeding was initially described by Zimmermann (1983) as “designed (i.e., predetermined) leaks in plant cell walls”, depending on the pore size and pressure difference between conduits. He suggested that embolism propagation occurs through pit membranes of axially adjacent conduits. Based on a theoretical example, a gas-liquid interface would be pulled across a pressure difference of 1.4 MPa from an air-filled conduit into a sap-filled one via a simple pore with a fixed diameter of 200 nm (Zimmermann, 1983; see Fig. 1 for a traditional illustration of air-seeding through smaller pores). The underlying idea behind air-seeding, however, is much older and can be traced back at least to the late 19^th^ century. Dixon and Joly (1894) suggested that the stability of xylem sap under negative pressure comes from internal stability of xylem sap, and from “the property of the pit-membranes to oppose the passage of free gas". Renner (1915) did not formulate an explicit air-seeding mechanism, but he recognized that xylem sap is in a physically metastable state under negative pressure, that the existence of continuous columns of xylem sap is difficult to confirm, and that large gas bubbles may expand to fill up conduits. Moreover, Oertli (1971) anticipated the physical principle underlying air-seeding by proposing that embolism occurs when air is aspirated into xylem conduits through pores in conduit walls, without suggesting the role of pit membranes in bordered pits.

**Fig. 1.**
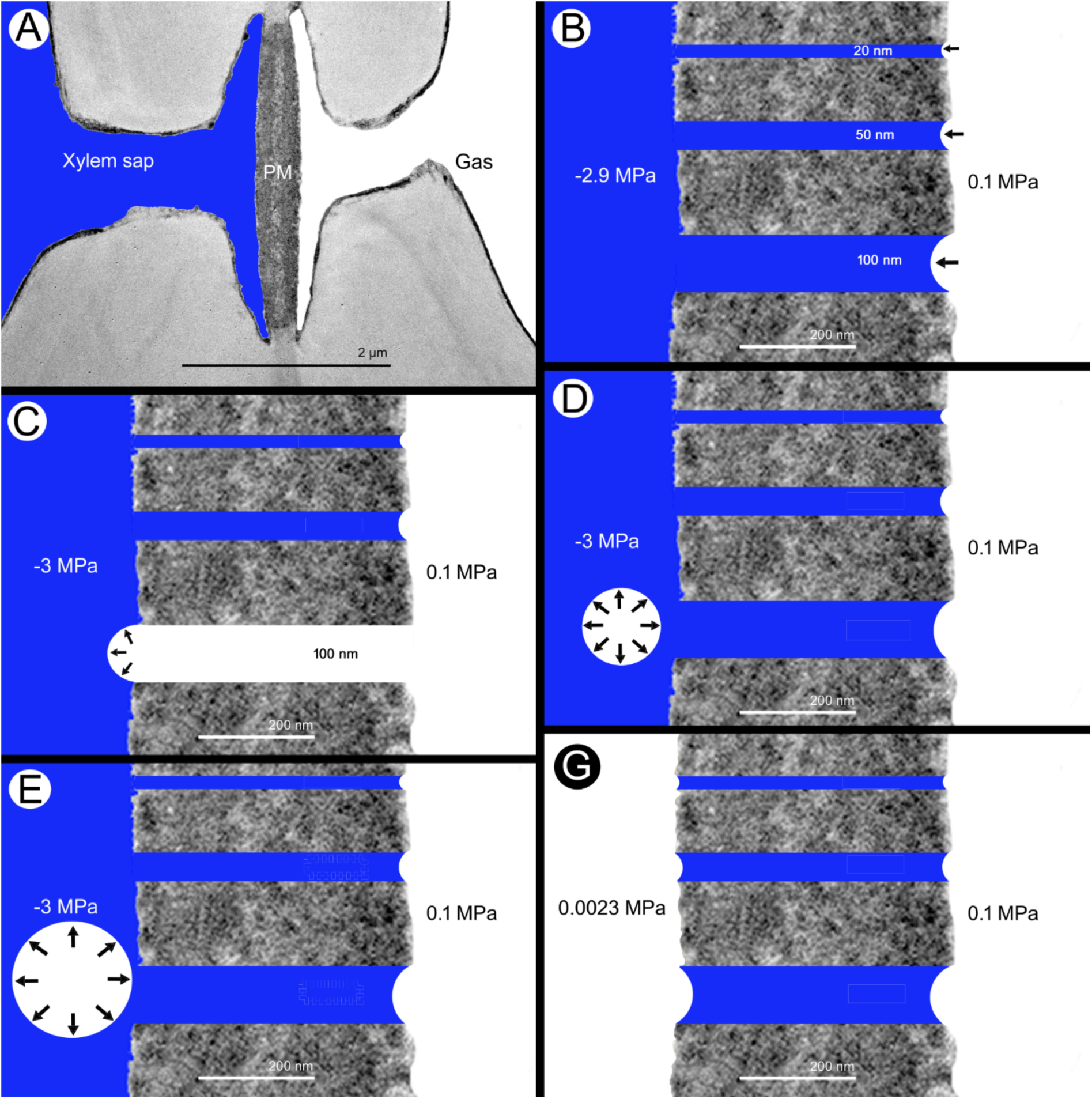
Illustration of the traditional air-seeding hypothesis showing gas-liquid interactions between an embolised and a sap-filled conduit (A), assuming a simplified pit membrane structure (B-F). **A.** An intervessel pit of *Citrus sinensis* (L.) Osbeck imaged with a transmission electron microscope. The left vessel is sap-filled and under negative pressure, while the right vessel is embolised. PM = pit membrane; the pit membrane thickness is not considered to affect air-seeding. **B.** Three pores are shown in a pit membrane; these are circular, simple, and have variable diameters (20, 50, and 100 nm). No polar lipids occur at gas-liquid interfaces. The gas-liquid meniscus seals off the pores, with the radius of the curvature being less than the radius of the pore. **C.** Assuming the surface tension of pure water, a meniscus moves across the largest, 100 nm pore as soon as a pressure difference of 2.9 MPa has been reached. The meniscus reaches the conduit lumen of the sap-filled conduit, where it grows rapidly. The 50 and 20 nm pores will remain sealed off until pressure differences of 5.8 and 14.5 MPa have been achieved, respectively. **D.** Based on Zimmermann (1983), the growing bubble may or may not detach from the wall and the pore reseals because the growing bubble volume (black arrows) causes briefly a local pressure increase. **E.** The non-stable bubble expands rapidly (black arrows), filling the entire vessel. **F.** The left vessel has become embolised, and is initially filled with water vapour (0.0023 MPa at 20°C), but the pressure will gradually increase to atmospheric pressure (0.1MPa; Wang et al., 2015a, b), similar to the embolised vessel on the right.

Numerous studies experimentally tested and supported Zimmermann’s air-seeding hypothesis, indicating that interconduit pit membranes represent the Achilles’ heel of xylem and play a major role in drought-induced embolism propagation (Lewis, 1988; Sperry and Tyree, 1988; Tyree and Sperry, 1989; Cochard *et al*., 1992; Hacke and Sperry, 2001). At the same time, pit membranes were also interpreted as efficient safety valves, preventing easy spreading of the gas phase, until a certain safety threshold or “air-seeding pressure” was achieved. While many studies supported air-seeding as a phenomenon that explains spatial embolism spread from one conduit to another (Brodersen *et al*., 2013; Choat *et al*., 2016; Wason *et al*., 2021), a new body of research creates the need to re-evaluate the mechanisms underlying embolism propagation.

The term air-seeding has become canonical in literature on plant water transport (Zimmermann, 1983; Sperry and Tyree, 1988; Tyree and Sperry, 1989). Many studies have treated air-seeding as a near-synonym of embolism propagation, framing it as a well-established mechanism underlying drought-induced embolism (Choat *et al*., 2008; Lens *et al*., 2013). However, recent findings on the relationship between pit structure and embolism propagation in xylem conduits make clear that the simplicity of the air-seeding model needs to be replaced by a much more complex set of biophysical processes and conditions (Rockwell *et al*., 2014; Jansen *et al*., 2018). Indeed, considerable progress has been made in quantifying embolism resistance (Cochard *et al*., 2013; Lamarque *et al*., 2018; Paligi *et al*., 2023), and in characterising vessel and bordered pit characteristics (Wheeler *et al*., 2005; Li *et al*., 2016; Kaack *et al*., 2019, 2021; Pereira *et al*., 2020; Levionnois *et al*., 2021). Moreover, new concepts have emerged regarding xylem sap physiochemistry, such as dynamic surface tension driven by polar, insoluble lipids (Scott *et al*., 1960; Esau, 1965; Schenk *et al*., 2017; Yang *et al*., 2020; Levionnois *et al*., 2022) surfactant-coated nanobubbles (Schenk *et al*., 2015, 2017; Ingram *et al*., 2021, 2023; Guan *et al*., 2022), and oversaturated concentrations of dissolved gas in xylem sap (Schenk *et al*., 2016; Marion *et al*., 2026).

Because various novel findings are incompatible with the assumptions underlying air-seeding, this paper provides a critical re-evaluation of the mechanisms of embolism propagation. We present an updated mechanistic framework for gas-liquid interactions in xylem conduits. Embolism propagation occurs in the xylem of all vascular plants, including ferns, gymnosperms, and angiosperms. Based on similarities in conduit and pit characteristics, embolism propagation in ferns is mechanistically likely similar to that in angiosperms (Suissa and Friedman, 2021; Pittermann *et al*., 2023). Torus-margo pit membranes in gymnosperm tracheids, however, may exhibit a different mode of embolism propagation (Bouche *et al*., 2014). Pit-membrane aspiration (i.e., complete sealing of the torus-bearing pit membrane against the outer pit aperture), for instance, is a common, well-established process in conifers with torus-margo pit membranes (Bailey, 1913; Beikircher *et al*., 2010; Zelinka *et al*., 2015), but not in angiosperms (Tixier *et al*., 2014; Carmesin *et al*., 2023). Because the dynamic nature of embolism propagation substantially increases the complexity of the subject, we limit the scope of this review to angiosperms, excluding other vascular plant groups.

### Terminology matters - words define mechanisms

Here, we provide a definition of the key processes and structures to avoid misinterpretations and confusion. A brief glossary of terms is also provided as supplementary information file (**SI1**).

**Air-seeding** has traditionally been defined as the mechanism by which embolism propagates from an embolised conduit containing gas at atmospheric pressure to a neighbouring sap-filled conduit through bulk flow of gas across pit membranes in bordered pits (**Fig. 1**; Zimmermann, 1983). The chemical composition of this gas is assumed to be similar to air, while the term seeding refers to the initial entry of a gas phase into a sap-filled conduit. Importantly, this entry of a tiny gas bubble in sap under negative pressure by capillary failure is assumed to grow directly into an embolism (see below for mechanistic details).

While there is valid support for spatial spreading of embolism between conduits (Brodribb *et al*., 2016; Choat *et al*., 2016; Korhonen *et al*., 2026), current mechanistic insights behind embolism propagation are not in line with air-seeding. Therefore, we use the term air-seeding as strictly referring to the traditional concept (Zimmermann, 1983; Sperry and Tyree, 1988; Cruiziat *et al*., 2002), and use the general term **embolism propagation** to describe the biophysical processes and conditions that underlie two different processes: bubble generation, and embolism formation. **Bubble generation** relates to the formation of nanobubbles, ranging between 20 to 300 nm in radius (Ingram *et al*., 2023), which takes place in pit membranes. **Embolism formation** is defined as the mechanistic process from expanding/coalescing nanobubbles in xylem sap to embolism of a conduit. Bubble generation is required for embolism formation, but bubble generation does not automatically lead to embolism. Both processes are interrelated to each other, but are temporarily and spatially separated. Unlike air-seeding, the entry of a single air bubble into a sap-filled conduit does not necessarily result in embolism (Schenk *et al*., 2015, 2017; Ingram *et al*., 2021).

We use the term **embolism** to describe the phase change of a sap-filled to a gas-filled conduit, which is known to block water transport and to occur at relatively high levels of drought stress. Initially, a recently embolised conduit includes 100% water vapour with a pressure close to vacuum, but will become filled with gas at atmospheric pressure, with a chemical composition that is likely similar to air (Wang *et al*., 2015a, b; Silva *et al*., 2024). The term **cavitation**, which is commonly defined as the nucleation of vapour bubbles in a liquid (Caupin and Herbert, 2006), is frequently misapplied in plant physiology to denote gas bubble expansion and often conflated with embolism (Zwieniecki and Secchi, 2015). Since xylem embolism is formed from expansion of a pre-existing gas bubble under decreasing pressure in a liquid, without nucleation, this process cannot be called cavitation.

**Conduits** in xylem tissue of angiosperms include both multicellular vessels and unicellular tracheids. Vessels are characterised by perforation plates between vessel elements, which are lacking in tracheids. Both conduit types, however, share bordered pits in their interconduit walls, which are also defined as end walls, even if the joint interconduit pit fields in these walls are located away from the ends of the vessels. Unlike interconduit pits, fibres show indistinctly bordered or simple pits between adjacent fibres (Sano *et al*., 2011; Olson, 2023). Interconduit pits include intervessel, intertracheid, and vessel-tracheid pits, but not vessel-parenchyma pits. The anatomical differences between xylem conduits in primary and secondary xylem can be large and may affect embolism resistance (Lens *et al*., 2022). However, the underlying mechanism of embolism propagation is largely similar in both xylem types and various plant organs. Therefore, we use the term xylem in a broad sense, including both primary and secondary xylem across various plant organs.

The main **porous medium characteristics** of pit membranes inside bordered pits include: (1) the porosity (i.e., the pore volume fraction), (2) the geodesic tortuosity (i.e., the ratio of the mean shortest flow path length to the pit membrane thickness), and (3) the constrictivity as indicator of **pore constrictions**, which are defined as a locally narrowed spaces within a three-dimensional pore pathway (for details and full definitions, see Zhang *et al*. (2020).

Interconduit pit membranes provide a **pore network**, defined as the three-dimensional, interconnected ensemble of nanoscale voids and solid matrix constrictions within a hydrated pit membrane. This network determines the structural connectivity of the membrane and underlies its hydraulic and gas transport properties. A single **pore** refers to a local void space between structural elements of the pit membrane through which water, solutes, or gases may pass depending on local boundary conditions. The pore network thus represents the full structural connectivity of the pit membrane, from which multiple potential connectivity routes can emerge under hydrated states. A **pore-pathway**, however, refers to a transient, hydraulically active percolation route through the pore network, composed of sequentially connected pore spaces and constrictions that become functionally continuous under specific pressure and interfacial conditions. Rather than representing a fixed channel, a pore pathway is a state-dependent realisation of connectivity within the pore network and may change dynamically (e.g., with water potential, surface tension effects, local presence of polar lipids and proteins, an electric double layer, and mechanical deformation of the pit membrane). The pore-pathway concept is particularly useful for modelling interfacial phenomena such as passage of gas-liquid interfaces across pit membranes and bubble generation.

The **pit border** represents the overhanging secondary cell wall, which encloses the pit chamber (Fig. 2). The **pit chamber** represents a fairly large (ca. 10-30 µm³), semi-enclosed volume surrounded by the roof of the pit border and the pit membrane. When protuberances from the secondary cell wall occur on the pit border or near the outer pit aperture, the bordered pits are vestured (Jansen *et al*., 2001).

**Fig. 2.**
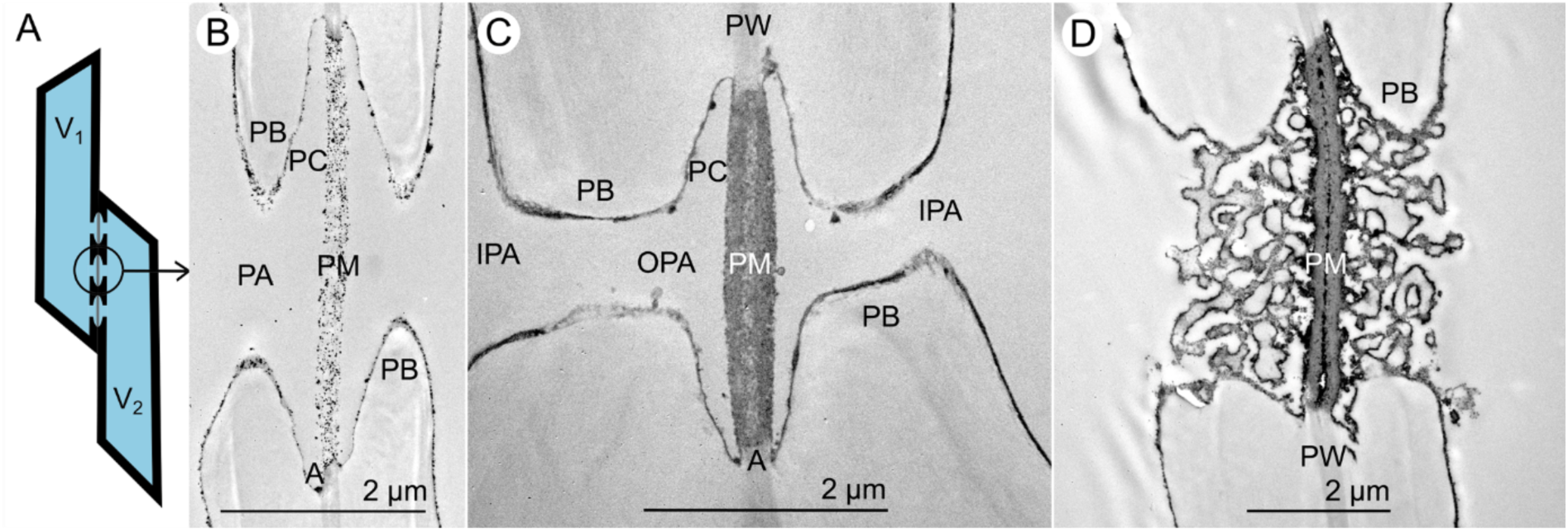
Micromorphology of bordered pits between neighbouring xylem conduits in angiosperms. Two neighbouring vessels with intervessels pits are shown in A. Images based on transmission electron microscopy represent *Corylus avellana* L. (B), *Citrus sinensis* (L.) Osbeck (C), and *Dryobalanops lanceolata* Burk (D). While a relatively thin secondary wall occurs in B, a thicker secondary wall distinguishes the inner pit aperture (IPA) from the outer pit aperture (OPA) in C. Vestures, which represent protuberances from the secondary cell wall, fill up the pit chambers in D. The vestured pit is characterised by large pit apertures and a relatively short pit border. The pit membrane (PM) represents the modified primary cell wall (PW), with a pit membrane annulus (A) near the edge of the pit border, and shows considerable variation in thickness between the species shown. The thin, dark coating on all conduit wall surfaces and pit membranes include polar lipids that become visible after treatment with OsO_4_. These polar, insoluble lipids affect the wettability and electrostatic interactions between charged interfaces. Structural details are indicated by acronyms at selected positions only; not all corresponding structures are labelled in B, C, and D. A = annulus, IPA = inner pit aperture, OPA = outer pit aperture, PA = pit aperture, PB = pit border, PC = pit chamber, PM = pit membrane, PW = primary wall.

### The air-seeding concept – before lipids re-entered the scene

The air-seeding concept as proposed by Zimmermann (1983) describes how a gas-liquid interface moves across the pit membrane of an interconduit pit and leads to embolism in a neighbouring conduit (Table 1; Fig. 1).

**Table 1:**
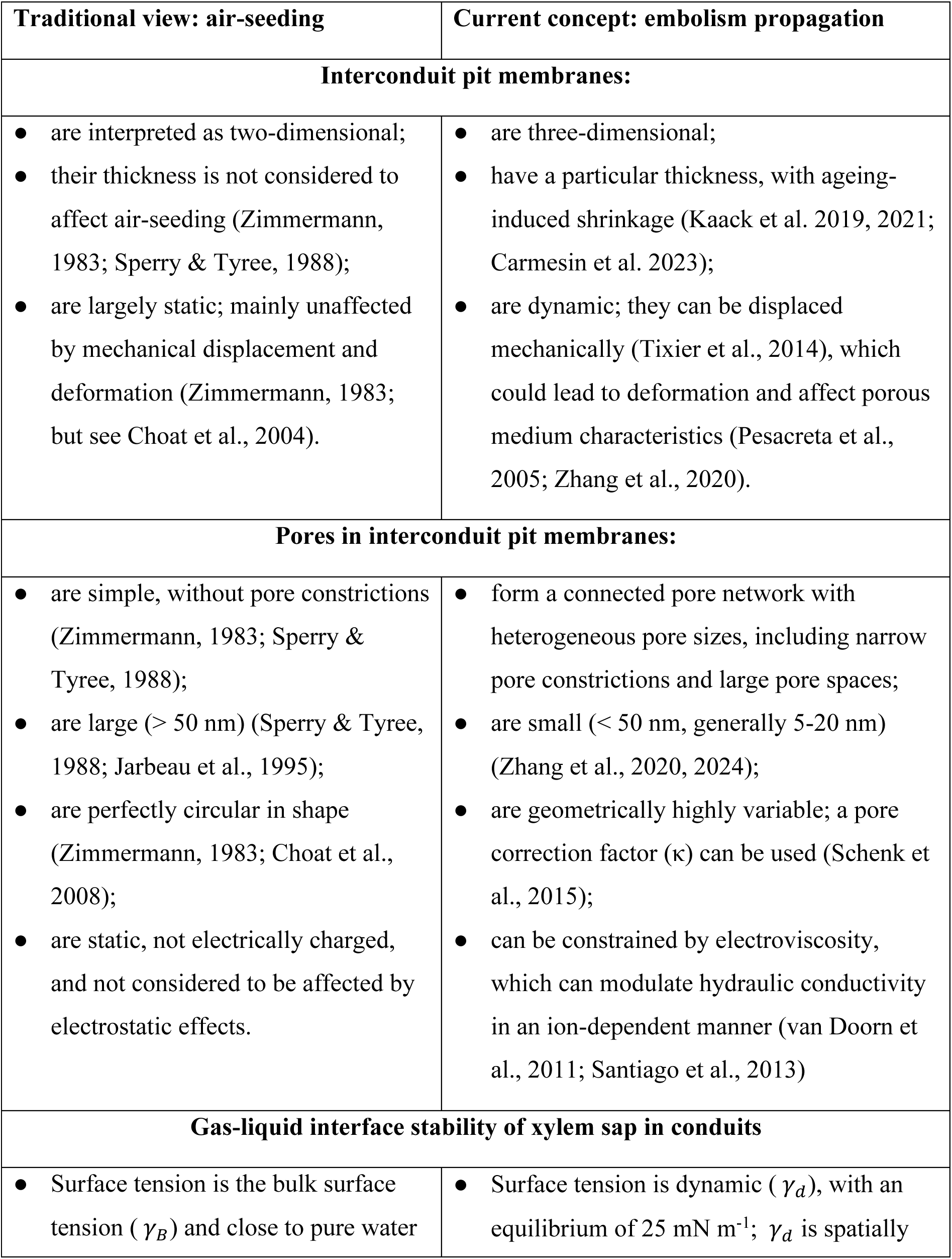
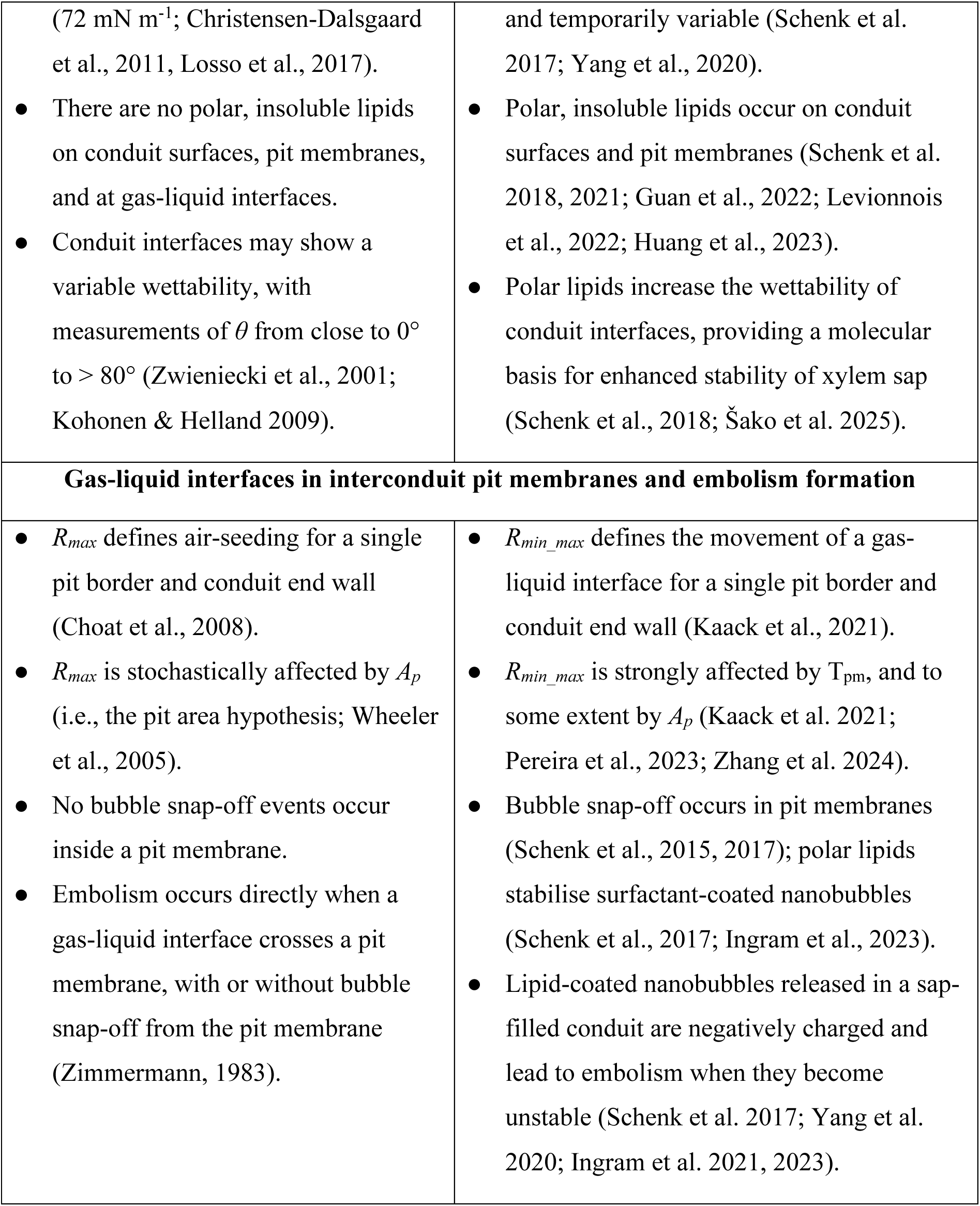
Summary and comparison of the traditional air-seeding concept (Fig. 1) and the current embolism propagation framework (Fig. 4). The related processes and biophysical concepts are arranged based on their location in xylem. Some key references are provided. For details, see the manuscript.

| Traditional view: air-seeding | Current concept: embolism propagation |
| --- | --- |
| **Interconduit pit membranes:** |  |
| • are interpreted as two-dimensional;<br>• their thickness is not considered to affect air-seeding (Zimmermann, 1983; Sperry & Tyree, 1988);<br>• are largely static; mainly unaffected by mechanical displacement and deformation (Zimmermann, 1983; but see Choat et al., 2004). | • are three-dimensional;<br>• have a particular thickness, with ageing-induced shrinkage (Kaack et al. 2019, 2021; Carmesin et al. 2023);<br>• are dynamic; they can be displaced mechanically (Tixier et al., 2014), which could lead to deformation and affect porous medium characteristics (Pesacreta et al., 2005; Zhang et al., 2020). |
| **Pores in interconduit pit membranes:** |  |
| • are simple, without pore constrictions (Zimmermann, 1983; Sperry & Tyree, 1988);<br>• are large ($> 50$ nm) (Sperry & Tyree, 1988; Jarbeau et al., 1995);<br>• are perfectly circular in shape (Zimmermann, 1983; Choat et al., 2008);<br>• are static, not electrically charged, and not considered to be affected by electrostatic effects. | • form a connected pore network with heterogeneous pore sizes, including narrow pore constrictions and large pore spaces;<br>• are small ($< 50$ nm, generally 5-20 nm) (Zhang et al., 2020, 2024);<br>• are geometrically highly variable; a pore correction factor ($\kappa$) can be used (Schenk et al., 2015);<br>• can be constrained by electroviscosity, which can modulate hydraulic conductivity in an ion-dependent manner (van Doorn et al., 2011; Santiago et al., 2013) |
| **Gas-liquid interface stability of xylem sap in conduits** |  |
| • Surface tension is the bulk surface tension ($\gamma_B$) and close to pure water | • Surface tension is dynamic ($\gamma_d$), with an equilibrium of $25 \text{ mN m}^{-1}$; $\gamma_d$ is spatially |
| (72 mN m<sup>-1</sup>; Christensen-Dalsgaard et al., 2011, Losso et al., 2017).<br>• There are no polar, insoluble lipids on conduit surfaces, pit membranes, and at gas-liquid interfaces.<br>• Conduit interfaces may show a variable wettability, with measurements of $\theta$ from close to 0° to > 80° (Zwieniecki et al., 2001; Kohonen & Helland 2009). | and temporarily variable (Schenk et al. 2017; Yang et al., 2020).<br>• Polar, insoluble lipids occur on conduit surfaces and pit membranes (Schenk et al. 2018, 2021; Guan et al., 2022; Levionnois et al., 2022; Huang et al., 2023).<br>• Polar lipids increase the wettability of conduit interfaces, providing a molecular basis for enhanced stability of xylem sap (Schenk et al., 2018; Šako et al. 2025). |
| **Gas-liquid interfaces in interconduit pit membranes and embolism formation** |  |
| • $R_{max}$ defines air-seeding for a single pit border and conduit end wall (Choat et al., 2008).<br>• $R_{max}$ is stochastically affected by $A_p$ (i.e., the pit area hypothesis; Wheeler et al., 2005).<br>• No bubble snap-off events occur inside a pit membrane.<br>• Embolism occurs directly when a gas-liquid interface crosses a pit membrane, with or without bubble snap-off from the pit membrane (Zimmermann, 1983). | • $R_{min\_max}$ defines the movement of a gas-liquid interface for a single pit border and conduit end wall (Kaack et al., 2021).<br>• $R_{min\_max}$ is strongly affected by $T_{pm}$, and to some extent by $A_p$ (Kaack et al. 2021; Pereira et al., 2023; Zhang et al. 2024).<br>• Bubble snap-off occurs in pit membranes (Schenk et al., 2015, 2017); polar lipids stabilise surfactant-coated nanobubbles (Schenk et al., 2017; Ingram et al., 2023).<br>• Lipid-coated nanobubbles released in a sap-filled conduit are negatively charged and lead to embolism when they become unstable (Schenk et al. 2017; Yang et al. 2020; Ingram et al. 2021, 2023). |

The spatial distribution of embolism propagation from an embolised conduit to a neighbouring, sap-filled conduit, has been confirmed experimentally. Novel imaging methods of dehydrating xylem samples, for instance, such as X-ray micro-computed tomography (microCT) observations (Brodersen *et al*., 2013; Choat *et al*., 2016; Wason *et al*., 2021) and the optical method (Brodribb *et al*., 2016) indicate that embolism almost never forms in a xylem conduit that is surrounded by sap-filled conduits only (Knipfer *et al*., 2015; Lens *et al*., 2022). Therefore, it can be assumed that *de novo* embolism formation is rare, although it cannot be fully excluded, and is hard to detect experimentally.

An important question, which has not been addressed in many studies, is where the first embolism comes from. The origin of the very first embolised conduit in a plant is likely associated with the development of xylem tissue, the type of xylem (primary vs secondary xylem), the functional longevity and mechanical properties of xylem conduits, the plant organ, and interconnectivity of conduits and vascular bundles along the entire xylem pathway from small roots to minor veins in leaves. Conduits in primary xylem, such as ring-tracheids or tracheary elements with a helical deposition of secondary wall, can be relatively vulnerable to mechanical stress, deformation, or damage, which also make them relatively vulnerable to embolism (Choat *et al*., 2005, 2016). Moreover, it is possible that air-entry from apoplastic spaces near the pith takes place across the walls of primary xylem conduits, suggesting that gas movement could be important to induce the first embolised conduit. Alternatively, gas-filled apoplastic spaces could also be produced by turgor loss of vessel-associated parenchyma cells (Tomasella *et al*., 2026). The difference with embolism propagation between conduits is that gas movement may not take place between neighbouring conduits, but between apoplastic space and a primary xylem conduit. Importantly, conduits are always interconnected to other conduits (Zimmermann, 1971; Burggraaf, 1972). Despite some degree of hydraulic compartmentalisation, conduits almost always overlap and practically never end in isolation (André, 2005). Therefore, perennial and woody plants are likely to have nearly always a certain amount of embolised, dysfunctional conduits in their xylem from where embolism could spread.

Quantitatively, the critical xylem pressure difference (*ΔP_air-seeding_)* that promotes the crossing of a gas-liquid interface across an end wall has been estimated by the Young-Laplace equation:

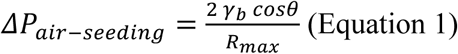

Here, *γ_b_* (N/m) represents the bulk surface tension of xylem sap, θ (°) is the liquid-solid contact angle with the pore wall, and *R*_max_ (nm) is the radius of the largest of all pores in an end wall. For an overview of all acronyms and definitions, see Table 2.

**Table 2:** Alphabetical list of acronyms used with their definition and units.

| Acronym | Definition | Comments | Units |
| --- | --- | --- | --- |
| $\beta$ | Pore shape correction factor | $0 \leq \beta \leq 1$ with values close to 1 for circular shapes, and much lower values for slit-like shapes (Emory 1989) | / |
| $\gamma_b$ | Bulk surface tension of xylem sap | $\gamma_b$ of xylem sap is typically close to the surface tension of pure water (72 mN m<sup>-1</sup>) or somewhat lower. | mN m<sup>-1</sup> |
| $\gamma_d(x, t)$ | Dynamic surface tension of xylem sap | $\gamma_d(x, t)$ is spatially and temporally variable, with values from 19 mN m<sup>-1</sup> to 68 mN m<sup>-1</sup> (Yang et al., 2020) | mN m<sup>-1</sup> |
| $\Delta P_{air-seeding}$ | The critical pressure to induce air-seeding | Can be estimated based on the Young-Laplace equation. | MPa |
| $\Delta P_{bubble}$ | The critical pressure for movement of a gas-liquid interface across a pit membrane | A modified Young-Laplace equation can be used to estimate $\Delta P_{bubble}$, integrating dynamic processes such as $\gamma_d, \beta, \theta, \Delta R_{mech}, \lambda_D$ | MPa |
| $\Delta R_{mech}$ | The change in pore radius due to pit membrane deformation | When the pit membrane is displaced by mechanical strain, the pore radius can become larger or smaller by stretching or bending of the pit membrane. | nm |
| $\epsilon_0$ | The vacuum permittivity | $\epsilon_0 = 8.854 \times 10^{-12} \text{ C}^2 \cdot \text{N}^{-1} \cdot \text{m}^{-2}$ | C<sup>2</sup>·N<sup>-1</sup>·m<sup>-2</sup> |
| $\epsilon$ | The relative permittivity or dielectric constant of water | $\epsilon$ for water at 20°C is 80 | / |
| $\theta$ | Contact angle of the gas-liquid interface with a conduit wall or pit membrane pore | For pit membranes, values between 10° and 60° are likely, and dynamically variable. | ° |
| $k$ | Pit membrane permeability index | Based on empirical data from Hacke et al. (2006), it can be estimated as $10^{-13}$ m<sup>2</sup> when using Darcy's law:<br>$k = \mu T_{pm} / R_m$<br>($R_m$ = pit membrane resistance) | m<sup>2</sup> |
| $\lambda_D$ | Debye length | The thickness of the electric double layer, which depends on the ionic concentration of xylem sap. This value is likely between 1 and 10 nm. | nm |
| $\mu$ | Viscosity of a fluid | For xylem sap, we assume $\mu = 0.001$ Pa s (at 20°C). | Pa s |
| $\nu$ | Poisson ratio | Can be estimated as 0.3 for cellulose-rich, soft material (Tixier et al., 2014). | / |
| $\chi$ | Pore deformation factor quantifying how pit membrane deflection is translated locally into changes in the effective radius of $R_{\min-\max}$ | $0 \leq \chi \leq 1$; this parameter links $w_{\max}$ with $\Delta R_{\text{mech}}$:<br>$\Delta R_{\text{mech}} = \chi w_{\max}$ | / |
| $\Pi_{el}(h)$ | The electrostatic disjoining pressure as a function of the height of a water film ($h$) that covers a pore constriction wall | Quantified in Equation S4 of Supplementary Information file 2 | Pa |
| $\sigma$ | The surface charge density | Expressed as C m<sup>-2</sup>, also as elementary charges per nm<sup>2</sup> (e/nm<sup>2</sup>); estimated values for pit membranes vary from 0.1 to 0.5 e nm<sup>-2</sup> (see SI). | C m<sup>-2</sup> |
| $A_p$ | The total intervessel pit membrane area of a vessel with average vessel dimensions | Can be modelled based on Wheeler et al. (2005), which requires vessel diameter, vessel length, intervessel contact fraction, and intervessel pit field fraction. | m<sup>2</sup> |
| $E$ | Young's modulus of the pit membrane | Estimated as 57 MPa for never-dried, fresh intervessel pit membranes of *Clematis vitalba* (Carmesin et al., 2023). | MPa |
| $h$ | Thickness of the water film on a charged surface, whose outer boundary is a water–vapour interface. | The thickness of the water film is proportional to the Debye length, and affected by the xylem sap pressure, with more negative pressure resulting in a thinner film. | m |
| $I$ | Ionic strength of xylem sap | Xylem sap ionic strength is relatively low and variable, with a typical estimation of roughly 1 to 10 mM under normal (i.e., non-stressed) conditions. | mol L<sup>-1</sup> |
| $P$ | Pressure of xylem sap | During transpiration, xylem sap is negative, with values from -0.1 to < -5MPa | MPa |
| $r_m$ | Pit membrane radius | Pit membranes can be assumed to be circular, and this parameter can be estimated as the equivalent circle radius in case pits are oval or elongated. | μm |
| $R_{eff}$ | The effective pore constriction diameter in a pit membrane. | The effective pore is dynamic, accounting for pressure-induced membrane deformation. | nm |
| $R_{max}$ | The largest pore in an infinitely thin pit membrane that is composed of a single layer. | This pore is assumed not to include a pore constriction. | nm |
| $R_{min-max}$ | The largest of the most minimal pore constrictions in a pore pathway. | $R_{min-max}$ can be estimated at the individual pit membrane level, or the vessel level based on a stochastic model presented in Kaack et al. (2021). | nm |
| $T_{pm}$ | Interconduit pit membrane thickness | Average pit membrane thickness as measured based on TEM in the centre. Measurements near the annulus and central position should be measured separately (Kaack et al., 2021). | nm |
| $W_{max}$ | Maximal central deflection of the pit membrane | Can be estimated with the Kirchhoff-Love plate theory, and a flexural rigidity ($D$) equation for pit membranes (Tixier et al., 2014); $D = \frac{E T_{pm}^3}{12 (1 - \nu)}$ | nm |

If we apply equation 1 to determine the critical pressure that induces embolism, which has been referred to as the air-seeding pressure, the following simplifying assumptions are made (Table 1; Zimmermann, 1983; Sperry and Tyree, 1988; Jarbeau, Ewers and Davis, 1995; Choat, Cobb and Jansen, 2008):

- Interconduit pit membranes consist of a single, homogenous layer. Simple pores of variable sizes directly cross the entire pit membrane, with a geodesic tortuosity equal to 1. These pores are perfectly circular, and show no pore constrictions (Fig. 1B-F). The pit membrane thickness and the pore network inside pit membranes are not considered to affect air-seeding (Zimmermann, 1983; Sperry and Tyree, 1988; Mrad *et al*., 2018, 2021).
- The radius of the largest pore in an interconduit end wall (*R*_max_) determines embolism spread (Jansen *et al*., 2009; Mrad *et al*., 2018, 2021). Also, the stochastic likelihood of having a high *R_max_* value in an end wall has been suggested to increase with the size of this end wall (Wheeler *et al*., 2005). The total intervessel pit membrane area per vessel (*A_p_*) has been linked to embolism resistance as the “pit area hypothesis”. Moreover, since pore sizes in pit membranes were generally found to be too small to match air-seeding pressures (Cronshaw, 1960; Murmanis and Chudnoff, 1979; Van Alfen *et al*., 1983; Jarbeau *et al*., 1995; Shane *et al*., 2000; Choat *et al*., 2003), the “rare pit hypothesis” has been proposed (Christman *et al*., 2009, 2012). This hypothesis, which proposes that embolism resistance is governed by the probability of encountering exceptionally large pore constrictions within a pit membrane population, should be conceptually distinguished from the pit area hypothesis.
- The bulk surface tension (*γ_b_*) is assumed to be similar or close to that of pure water (i.e., 72 mN m^-1^ at 20°C). Bulk surface tension measurements of xylem sap have shown high surface tensions, with variation from ca. 50 to 70 mN m^-1^ (Christensen-Dalsgaard *et al*., 2011; Losso *et al*., 2017).
- The liquid-solid contact angle (*θ*) is close to 0°, suggesting that xylem inner walls and pit membranes are highly hydrophilic, with a uniform, maximum wettability. However, experimental work contradicted this high wettability, and contact angles from close to 0° to > 80° have been measured (Zwieniecki and Holbrook, 2000; Kohonen, 2006; Kohonen and Helland, 2009; Brodersen *et al*., 2010). An increased contact angle would reduce the air-seeding pressure. McCully *et al*. (2014) suggested that bordered pits in vessels of maize roots have a dual nature: hydrophobic when embolised, but hydrophilic when wetted by sap.
- Any bubble or liquid-gas interface passing a pit membrane leads to a new embolised conduit, with an initial pressure close to water vapour pressure (0.0023 MPa at 20°C). The likelihood of embolism formation from a bubble seed is not determined by the bubble size, because any bubble would trigger a phase change from the metastable liquid under negative pressure to a void and form embolism. The lack of pore constrictions in pit membranes implies that bubble snap-off events do not occur. Zimmermann (1983), however, suggested that enlarging bubbles that enter a sap-filled pit border may or may not detach from the pit membrane, with a possibility that the pore can instantaneously reseal with water after admitting a bubble (Fig. 1E).
- Pit membrane displacement and changes in the pore size, which could be caused by a pressure difference across a bordered pit pair, are not considered. Porous medium characteristics of pit membranes, such as the pit membrane pore size, are assumed to be static, and not affected by displacement of cellulose microfibril aggregates. Likewise, the bordered pit geometry is not considered to affect air-seeding, although vestured pits have been suggested to support deflection and rupture of the pit membrane by a pressure difference across a bordered pit pair (Zweypfenning, 1978; Choat *et al*., 2004).

Overall, the air-seeding model gained widespread acceptance because of its attractive simplicity. There is strong experimental support that interconduit pits play a major role in vulnerability to embolism, and that embolism proceeds from an embolised to a sap-filled conduit (Sperry *et al*., 1996). However, the assumptions underlying Equation 1 and listed above have been challenged, as further discussed below.

### Novel insights challenging the air-seeding concept

Novel findings obtained over the last 15 years have substantially changed our understanding of embolism propagation (Table 1). These can be summarised into four key subjects: (1) the ultrastructure of pit membranes as mesoporous media, their chemical and mechanical properties (Kaack *et al*., 2021; Carmesin *et al*., 2023), (2) the concept of dynamic surface tension of xylem sap by polar, insoluble lipids (Schenk *et al*., 2017, 2021), (3) the generation and stability of surfactant-coated nanobubbles (Schenk *et al*., 2017; Guan *et al*., 2022; Ingram *et al*., 2023), and (4) an electric double layer, with pit membranes as electrochemical interfaces (van Doorn *et al*., 2011; Santiago *et al*., 2013; Ghildiyal *et al*., 2025). Here, we describe these key findings that challenge the traditional air-seeding concept, while mechanistic details about an updated framework of embolism propagation is provided in the next section.

#### (1) Interconduit pit membranes are three-dimensional, porous media

(Kaack *et al*., 2019; Zhang *et al*., 2020, 2024). They are mainly composed of cellulose microfibrils, which are typically grouped in microfibrillar aggregates with a diameter between 20 to 30 nm (Kaack *et al*., 2019). Pores in these pit membranes are ca. 5 to 50 nm in size, which is characteristic of mesoporous media (Fig. 3; Choat *et al*., 2004, 2008; Zhang *et al*., 2020, 2024). The thickness of pit membranes varies from ca. 200 nm to 1,000 nm (Li *et al*., 2016), and requires for accurate measurements the use of fresh samples based on transmission electron microscopy because xylem dehydration may lead to more than 50% shrinkage of pit membranes (Zhang *et al*. 2017, 2020). Porosity of pit membranes is very high (from 77% to 84%), geodesic tortuosity is close to 1 (between 1.02 and 1.03), and constrictivity is relatively high (from 0.60 to 0.81), indicating that multiple pore constrictions occur within each individual pore pathway (Kaack *et al*., 2019, 2021; Zhang *et al*., 2020). Unlike constrictivity, the porosity and geodesic tortuosity are likely independent of pit membrane thickness. Considering that pit membranes are multi-layered structures, the number of layers in a pit membrane does not appear to affect the density or the size of the cellulose microfibrillar aggregates. Understanding the porous medium characteristics of pit membranes enables us to apply Darcy’s law to pit membranes, which also provides an approach to estimate the pit membrane resistance to sap flow. Although measured values of the permeability index (*κ*) of pit membranes are unknown, this parameter has been estimated to be around 10^-13^ m^2^ (Li *et al*., 2020; Pereira *et al*., 2023; Table 2).

**Fig. 3.**
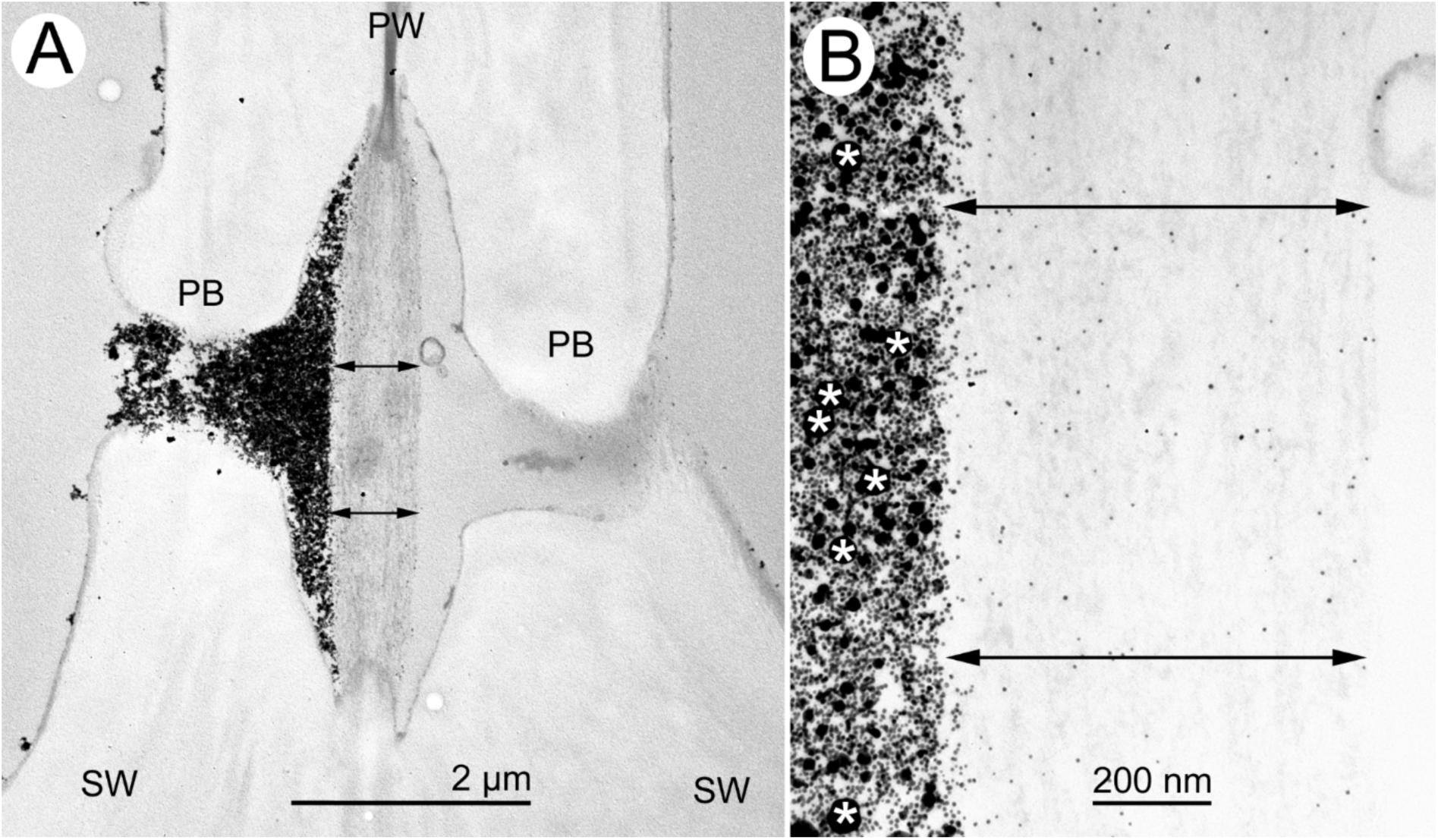
Intervessel pits of *Cinnamomum camphora* (L.) J.Presl not treated with OsO_4_ and injected with colloidal gold particles of four different diameters. Particle injection at a pressure of 200 kPa was done in a direction from the left to the right pit border, resulting in a dark, granular mass filling up the pit aperture and pit chamber (Zhang et al., 2024). Fig. B is a close up of the central pit membrane area in A. Only 5 and 10 nm particles penetrated the multi-layered pit membrane (outline indicated by double-sided arrows), while larger particles of 20 and 50 nm were not entering. Since OsO_4_ was omitted during the preparation of this sample, the polar, insoluble lipids lining the pit border and pit membrane are electron transparent and largely unnoticeable, contrary to the OsO_4_ treated samples in Fig. 2. The omission of OsO_4_ resulted in a low contrast of the pit membrane against the background, but increased the visibility of the colloidal gold particles. SW = secondary wall; PB = pit border, PW = primary wall, * = 50 nm particles

**Fig. 4.**
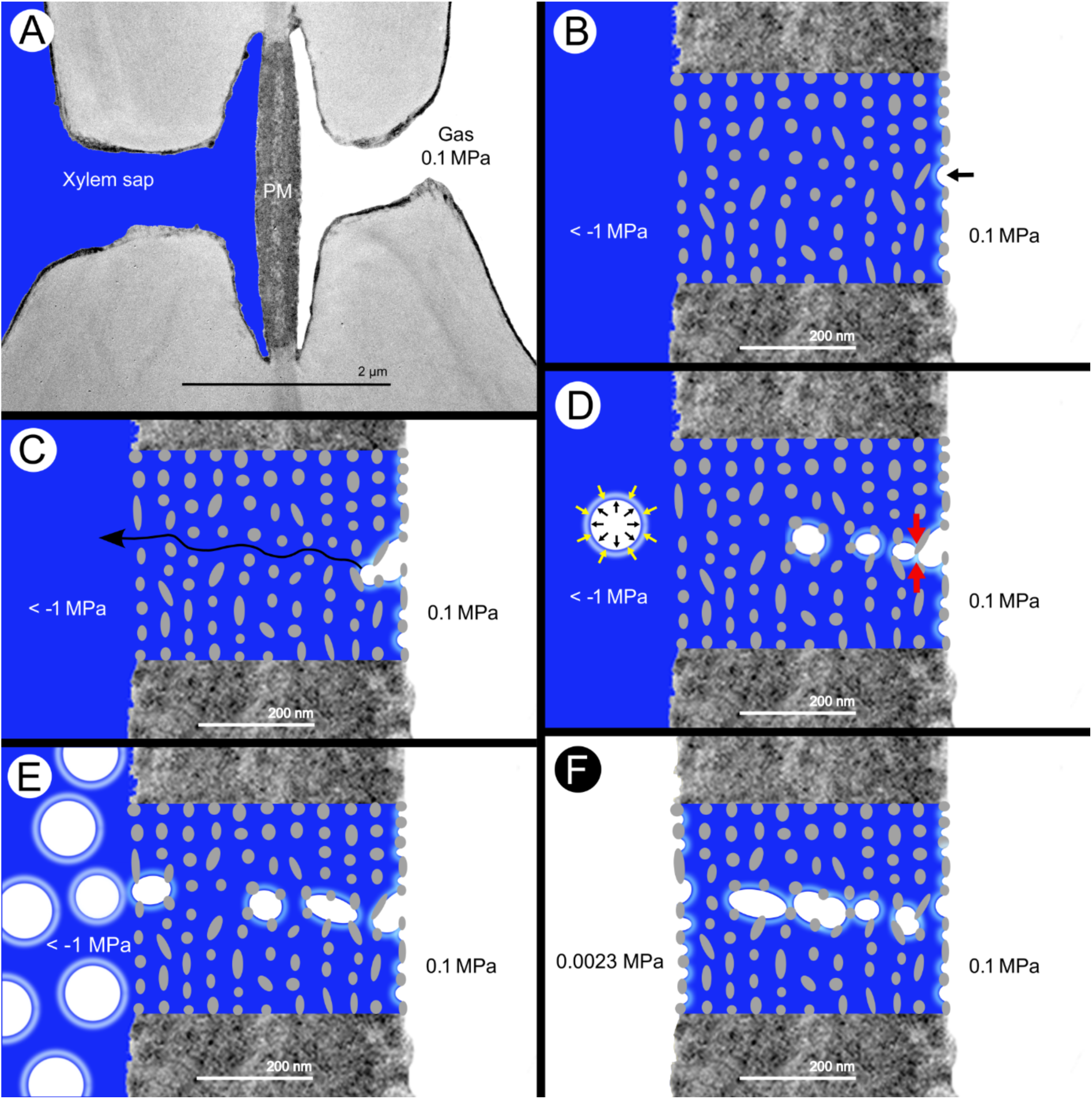
Gas-liquid interactions between an embolised and a sap-filled conduit (A, B, C), with bubble generation (D-E) and embolism formation (F) as two separate processes, emphasizing the dynamic processes of multiphase interactions with polar lipids as surfactants. The time scale between the depicted processes can be highly variable, ranging from less than milliseconds to many hours. **A.** An intervessel pit of *Citrus sinensis* (L.) Osbeck imaged with a transmission electron microscope. The left vessel is sap-filled and under negative pressure, while the right vessel is embolised. PM = pit membrane. Unlike Fig. 1A, the thickness of the pit membrane affects embolism propagation. **B.** Cellulose microfibril aggregates with a diameter of ca. 20 nm make up a pit membrane with a thickness of ca. 470 nm. This pit membrane includes 11 layers, has a porosity of ca. 80%, and a distance between microfibril aggregates of ca. 20 nm. The shape of the microfibril aggregates is circular, oval, or elongated, depending on their random orientation. The dynamic surface tension by polar lipids at the gas-liquid interface is represented by a luminous glow. The curvature of the gas-liquid meniscus can be variable, and generally somewhat higher than in Fig. 1B due to its reduced, dynamic surface tension. The black arrow indicates the largest gas-liquid meniscus. **C.** Depending on the local concentration of polar lipids at the gas-liquid interface and the dimensions of pore voids between the cellulose microfibril aggregates, gas-liquid menisci will move into the pit membrane. For reasons of simplicity, interactions along a single pore pathway are shown only, as shown by the black arrow crossing the entire pit membrane. The invading gas does not move as a simple gas-liquid front across the multi-layered pit membrane with pore constrictions. **D.** Snap-off events or Haines jumps (red arrows) occur when the radius of a pore constriction is less than half the radius of the pore void behind it. Bubble snap-off is driven by an increase of the local liquid pressure (due to bubble entry and low liquid compressibility), while minimalization of the surface area of a small, single bubble is thermodynamically favoured over large gas-liquid surface areas in a continuous, uninterrupted interface along the entire pore pathway. A bubble in a pit border of a sap-filled conduit can be stable due to counteracting forces on the bubble: internal gas pressure and a pulling force by the negative pressure of xylem sap cause expansion (black arrows), while the Laplace pressure pushes on the surface (yellow arrows). **E.** Snap-off events produce surfactant-coated nanobubbles that are inside the pit membrane and released in the neighbouring pit chamber. The lipids contribute to the stability of nanobubbles, which are negatively charged. Bubble propagation does not automatically lead to direct embolism formation because nanobubbles may remain stable or collapse. **F.** Nanobubbles can become unstable at certain yet unknown thresholds and cause embolism, depending on the concentration and nature of polar lipids, potential changes in temperature and pressure, and the concentration of dissolved gas in xylem sap. The left vessel became embolised and is filled with water vapour after nanobubbles became unstable by a combination of factors. The initial pressure of water vapour (0.0023 MPa at 20°C), will gradually increase to atmospheric pressure (0.1MPa; Wang et al., 2015a, b), as shown in the embolised vessel on the right.

Contrary to the air-seeding concept, which is based on *R_max_* in pit membranes assuming simple pores and no pore constrictions, movement of a fluid or gas-liquid interface in a pore pathway with multiple pore constrictions depends on the diameter of the smallest pore constriction along this path. For transport at the scale of an entire pit membrane, or even all interconduit pits in an end wall, we consider the largest of all minimal pore constrictions (*R_min-max_*) as representing the main bottleneck (Kaack *et al*., 2021). In other words, relevant here is the largest of all pore pathway minima. Assuming a pit membrane has only 100 pore pathways, with the narrowest pore constriction in all pore pathways varying from 5 nm to 18 nm, then *R_min-max_* is 18 nm, which is the largest of all pore pathway bottlenecks in that pit membrane. *R_min-max_* represents thus the most permissive pore pathway for gas-liquid interface passage and therefore the anatomically most relevant constriction governing transport. Unlike *R_max_*, *R_min-max_* explicitly incorporates the three-dimensional, multi-layered nature of pit membranes and avoids the unrealistic assumption of a single cylindrical pore through an infinitely thin membrane.

The concept of *R_min-max_* provides a mechanistic explanation for the relationship between pit membrane thickness and embolism resistance (Jansen *et al*., 2009; Lens *et al*., 2011; Plavcová *et al*., 2011; Li *et al*., 2016; Kaack *et al*., 2021; Levionnois *et al*., 2021; Isasa *et al*., 2023; Miranda *et al*., 2024). If pit membranes are thick, there is an increased number of pit microfibril layers, and pore constrictions. A high number of pore constrictions increases the likelihood that *R_min-max_* values, which can be stochastically modelled based on Kaack et al. (2021), become considerably smaller. Low values of *R_min-max_*, which are associated with high pit membrane thickness, would reduce gas-liquid interfaces and bubble generation, and increase embolism resistance. Considering that pore pathways traverse multiple layers of microfibril aggregates, high pit membrane thickness increases the number of constrictions encountered along each pore pathway. Consequently, both the mean and variance of *R_min-max_* are expected to decrease with increasing pit membrane thickness, making highly permeable pathways exponentially rare in thick pit membranes. This three-dimensional pit membrane structure and the number of pore constrictions encountered along a pore pathway essentially contradict the rare pit hypothesis, which relies on an effectively two-dimensional pit membrane concept (Christman *et al*., 2009, 2012; Li *et al*., 2020; Kaack *et al*., 2021). According to extreme-value statistics, the probability that all constrictions along a pathway remain large decreases rapidly as the number of constrictions increases, because every constriction represents another opportunity for a small bottleneck to occur. Therefore, the probability of large effective pore diameters becomes extremely low with each additional pore constriction along the path.

Moreover, interconduit pit membranes have been observed to shrink over time *in planta* (Schmid and Machado, 1968), which was found to be associated with increased embolism resistance in grapevine (Sorek *et al*., 2021). It is likely that this shrinkage occurs due to mechanical strain by flow, a pressure-difference across a sap-filled and embolised conduit, or due to ageing (e.g., transition from sapwood to heartwood). Shrinkage of interconduit pit membranes appears to be irreversible (Zhang *et al*., 2017, 2020). Clearly, more research is needed to determine how ageing and mechanical stress affect pit membrane ultrastructure and deformation, and how these changes influence sap transport, embolism resistance, and the functional life-span of sapwood (Sperry *et al*., 1991; Tixier *et al*., 2014; Hillabrand *et al*., 2016; Carmesin *et al*., 2023).

#### (2) Xylem sap has a dynamic surface tension at gas-liquid interfaces,

depending on the local concentration of polar lipids per surface area and temporal changes of these concentrations (Schenk *et al*., 2017; Yang *et al*., 2020). This dynamic surface tension (*γ_d_*) is fundamentally different from the equilibrium surface tension of bulk xylem sap (*γ_b_*), sometimes assumed to be around 72 mN m^-1^ for pure water at room temperature. The polar, insoluble lipids can strongly modify the surface tension of xylem sap, varying from 19 mN m^-1^ to 68 mN m^-1^, depending on the concentration of lipids at the surface (Yang *et al*., 2020). Molecular dynamics simulations have further revealed that both the temperature (Ingram *et al*., 2024) and water potential (Ingram *et al*., 2021) affect *γ_d_* at a given area per lipid. Nevertheless, low surface tensions are compatible with the range of embolism resistance observed in plants, especially when considering the smaller than previously assumed pore constrictions in pit membranes (Kaack *et al*., 2019; Zhang *et al*., 2024). Lower values of *γ_d_* also reduce the energy required to expand the gas-liquid interfaces, and therefore strongly affect the interfacial behaviour in pit membranes.

The dynamic surface tension is caused by polar, amphiphilic lipids in xylem sap (Schenk *et al*., 2017; Yang *et al*., 2020). Careful chemical analyses that control for cytoplasmic contamination from living xylem cells have shown that these lipids include galactolipids and phospholipids. These polar lipids were shown to be remnants of the cell membrane, cytoplasm, and plastids of living vessel elements, with plastids being the source of galactolipids (Dörmann and Benning, 2002). They are not soluble (Scott *et al*., 1960; Esau, 1965; Esau *et al*., 1966) and coat inner conduit walls and interconduit pit membranes (Schenk *et al*., 2017, 2018, 2021). The lipids are visible as a thin, electron-dense layer under a transmission electron microscope after treatment with osmium tetroxide (OsO_4_), a fixative and a stain, which binds to double carbon bonds in unsaturated fatty acid chains (Riemersma, 1968). The lipid layer is clearly visible in Fig. 2, but lacking in Fig. 3, which was not treated with OsO_4_. Since lipid micelles (i.e., small spherical aggregates of amphiphilic lipid molecules in xylem sap) are likely transported with xylem sap within a single vessel, they accumulate on interconduit pit membranes. When OsO_4_ is omitted in TEM fixation, pit membranes are largely invisible under TEM (Fig. 3). Also, due to an increasing amount of lipid deposition, pit membranes become gradually darker with ageing (Schmid and Machado, 1968). The lipid micelles, however, generally do not pass interconduit pit membranes, as shown experimentally by reduced lipid concentrations when consecutive amounts of sap are extracted from cut-open stems (Guan *et al*., 2022).

#### (3) Bubble generation by snap-off events in pit membranes leads to stable, surfactant-coated nanobubbles in xylem sap

Combined with the geometrically highly variable pore constrictions and pore voids, the potentially very low surface tension should yield bubble snap-off processes, which are also known as Haines jumps in porous media. They take place when a meniscus reaches a pore constriction and the capillary pressure becomes unstable, causing the interface to jump to the next constriction. Energetically more stable spherical bubbles can then be formed. For mechanistic details of snap-off events, see Schenk *et al*. (2017, 2021) and Ingram *et al*. (2023).

Studying these nanobubbles in the xylem of intact plants and under negative pressure is highly challenging. They can be quantified based on nanoparticle tracking analysis in extracted xylem sap, or imaged using freeze-fracture TEM (Schenk *et al*., 2017; Guan *et al*., 2022; Ingram *et al*., 2023; Huang *et al*., 2024). The radius of nanobubbles in extracted sap that is under atmospheric pressure is found to be between 20 and 300 nm, although the size of nanobubbles must be different in sap under negative pressure. Once bubbles are formed, their polar lipid coatings stabilise them, preventing expansion or contraction under highly negative or positive pressures, respectively (Dockar *et al*., 2019; Kanduč *et al*., 2020; Mi *et al*., 2025). Nanobubbles are filtered out from sap by interconduit pit membranes, as shown by consecutive sap extraction of stem samples, indicating that they are unlikely to pass between end walls of conduits (Guan *et al*., 2022). This means that nanobubbles can only be extracted from xylem sap in cut-open conduits of stem samples, but cannot be extracted from xylem sap in intact conduits. The reason is that the pores in interconduit pit membranes are too small (< 50 nm) to allow any nanobubbles through at their equilibrium size.

#### (4) Electric double layers occur at conduit walls and pit membranes

Electric double layers arise when charged surfaces interact with an electrolyte. In xylem, negatively charged surfaces occur on the hydrophilic head groups of amphiphilic polar lipids coating conduit walls and at least the outer surfaces of interconduit pit membranes, as well as on ionisable functional groups associated with pit membranes (Kaack *et al*., 2019). These charged surfaces attract oppositely charged ions (i.e., counter-ions) from the xylem sap, resulting in the formation of an electric double layer and an electrostatic force arising from this charge distribution. Consequently, in addition to capillary and dynamic surface tension forces, electrostatic interactions contribute to the physical environment in which water and gas move through bordered pits (van Doorn *et al*., 2011; Santiago *et al*., 2013; Ghildiyal *et al*., 2025).

The properties of the electric double layer are governed by two complementary components. First, the surface chemistry determines the surface charge density, zeta potential, and electroviscous behaviour, and is influenced by factors including lipid composition, lipid density, pH, and the chemical composition of the pit membrane. Second, the characteristic thickness of the electric double layer, expressed as the Debye length (*λ_D_*), is primarily determined by the electrolyte properties of the xylem sap, especially its ionic strength and, to a lesser extent, temperature. Thus, the charged surface establishes the electric potential, whereas the ionic composition of the xylem sap determines the distance over which this potential decays.

The hydraulic consequences of changing ionic strength are apparent as an "ionic effect", alterations in hydraulic conductance caused by changes in ionic composition of sap. This effect was assumed to result from swelling or shrinking of pectins in pit membranes (Zwieniecki *et al*., 2001; Nardini *et al*., 2011). However, this interpretation is hard to reconcile with evidence demonstrating that mature interconduit pit membranes contain no pectins, except within the annulus (i.e., very close to the pit border) (Plavcová and Hacke, 2011; Kim and Daniel, 2013; Herbette *et al*., 2015; Klepsch *et al*., 2016; Kaack *et al*., 2019). Most likely, changes in ionic strength modify the Debye length itself, thereby altering electrohydrodynamic interactions within nanometre-sized pore constrictions without requiring structural changes to the pit membrane (van Doorn *et al*., 2011; Santiago *et al*., 2013).

For typical xylem sap ionic strengths of approximately 5 to 20 mM (Marschner, 1995; Herdel *et al*., 2001; van Ieperen and van Gelder, 2006), the Debye length is expected to range between roughly 2 and 5 nm. If, for example, a pore constriction has a diameter of approximately 10 nm and we assume that this pore is bounded by two negatively charged walls, a Debye length of 4.3 nm (corresponding to an ionic strength of about 5 mM) would leave only about 1.4 nm of bulk-like fluid in the centre of the pore. Under these conditions, the electric double layers from opposite walls may substantially overlap, intensifying electroviscous drag on flow through the pore (Santiago *et al*., 2013). Consequently, the influence of electrostatic forces is expected to become most pronounced when pore dimensions approach approximately twice the Debye length.

Even if polar lipids may only be present on the outermost layers of pit membranes, the occurrence of electric double layers has several important implications for transport across pit membranes. First, pit membranes should be regarded as electrochemical interfaces rather than inert porous structures. Second, electrostatic interactions generate electroviscous drag on ion and water transport through pit membranes, thereby influencing permeability, hydraulic resistance, and local flow fields. Third, because electrohydrodynamic interactions occur simultaneously with capillary forces, dynamic surface tension, and multiphase flow, gas-liquid interfaces within pit membranes cannot be considered simple air-water menisci moving through static pores. Thus, these electrochemical processes, which arise from the coating of polar lipids, support a more dynamic view of embolism propagation in which bubble generation, nanobubble stabilization, and embolism formation represent distinct physical processes rather than a single air-seeding event.

##### Embolism propagation based on an updated gas-liquid interface concept

The new insights provided above largely contradict the simplifying assumptions of the air-seeding concept as a mechanistic understanding of embolism propagation between conduits, as helpful as it was. Indeed, the available evidence emphasizes that gas-liquid interactions in pit membranes are more complicated and dynamic than previously suggested. This complexity is mainly caused by the interactions of multiple phases, including gas, liquid xylem sap, solid material (mainly cellulose microfibril aggregates of pit membranes), and polar, insoluble lipids. The processes underlying these interactions are not static but dynamic, which implies that time also plays a role for the mechanisms driving embolism propagation. In agreement hereon, older xylem has been found most vulnerable to embolism propagation (Choat *et al*., 2005, 2016; Fukuda *et al*., 2015; Meixner *et al*., 2020; Weithmann *et al*., 2022). These time-dependent processes include dynamic changes in surface tension (e.g., surfactant rearrangement) and the mechanical properties of pit membranes, such as pit membrane deformation by displacement, and potential changes in porous medium characteristics (e.g., dynamic shifts in pore constrictions).

The new insights provided convincing evidence that gas-liquid interactions in pit membranes determine bubble generation. This process, however, is separate from embolism formation because nanobubbles, formed by snap-off events in foam-producing pit membranes, can be stable even under negative pressure (Ingram *et al*., 2023). The distinction between bubble generation and embolism formation requires separate treatments of the two phenomena, which are almost certainly causally related to each other. Bubble generation arises from the instability of gas-liquid interfaces within multi-layered pit membranes, where dynamic surface tension, pore network geometry, pressure-induced pit membrane deformation, and electrostatic forces play roles. Other factors, such as large and rapid changes in pressure, temperature, or nanobubble concentration, may then collectively determine the threshold for bubble expansion and therefore embolism formation.

To address the movement of a gas-liquid interface in a pit membrane, we provide here a conceptual first step towards such integration, although capturing these complex dynamics into a single, modified version of equation 1 remains a challenge. We define the bubble pressure difference (*ΔP_bubble_*) as the pressure required to move a gas-liquid-surfactant interface across a single pit membrane, or across all interconduit pit membranes in an end wall. Unlike the classical air-seeding pressure (*ΔP_air-seeding_*, Equation 1), *ΔP_bubble_* incorporates three departures from this idealised single-pore model: dynamic, rather than fixed, surface tension of xylem sap; irregular, multi-constriction pore geometry, including time-dependent membrane deformation; and an electrostatic disjoining pressure. Embolism resistance is therefore not a simple function of pore size, but an emergent property of coupled structural, electrochemical, hydraulic, and mechanical processes.

Formally, *ΔP_bubble_* follows a modified Young-Laplace equation, in which the capillary term is adjusted for multi-constriction geometry and membrane deformation, and augmented by an electrostatic disjoining pressure term (Hsu et al., 2003):

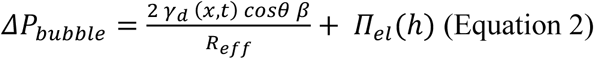

***γ_d_* (*x*, *t*) = dynamic surface tension,** with an equilibrium surface tension (*γ_e_*) around 25 mN m^-1^ (Yang *et al*., 2020). This parameter is not static but varies in space (x) and time (t), and could also be species-specific, depending on the composition and concentration of polar lipids (Schenk *et al*., 2021).

##### *Θ* = the liquid-solid contact angle with the pore wall

Although polar lipids are expected to accumulate on inner walls of xylem conduits and on pit membranes, their presence does not imply complete wetting (i.e., *θ* = 0°). Instead, the contact angle is likely finite and dynamically variable, reflecting the heterogeneous and non-equilibrium nature of lipid-coated interfaces on and possibly in pit membranes. With values between 10° and 60°, this factor could reduce *ΔP_bubble_* by half, which shows its large effect. We assume that no explicit time-dependence is needed for *Θ* because surfactant adsorption/desorption kinetics are likely much slower than the capillary-inertial time scale of snap-off itself.

***β* = a pore shape correction factor** (Emory, 1989) can be introduced to account for deviations in interfacial curvature because pore constriction shapes are geometrically highly variable and not perfectly circular. Values of *β* are between 1 and 0, with values close to 1 for perfect circular shapes, and much lower values for slit-like, elongated shapes. Estimating *β* based on imaging would be highly challenging due to the nanosized scale and potential deformation of pit membranes during sample preparation. Therefore, a sensitivity test for *β* varying between 0 to 1 is needed for evaluating its role in *ΔP_bubble_*.

***Π_el_*(*h*) = the electrostatic disjoining pressure** as a function of the height of the water film (*h*, expressed in m) that covers a pore constriction wall. For bulk flow through a pore constriction with a negatively charged surface, such as negatively charged cellulose surfaces with or without a negatively charged lipid coating, electroviscous resistance arises from an electroosmotically driven flow that opposes the pressure-driven flow (Santiago *et al*., 2013). Since lipid headgroup composition determines the surface charge density (*σ*) of the pore constriction wall, and the magnitude of *h* depends on *σ* together with the ionic concentration and pH of xylem sap, lipid coating plays a significant role in setting the disjoining pressure that resists bubble movement through pore constrictions. Amphiphilic lipids can strongly influence the electrochemical boundary conditions at pore surfaces, including surface charge and ion distribution, thereby modulating the effective impact of electric double layers on pore-scale transport. If polar lipids only coat the outermost layer of pit membranes (Schenk *et al*., 2018), electric double layers would be most pronounced at the pit membrane entrance, and electrostatic charges would be spatially heterogeneous. Thus, the effect of electric double layers can modulate hydraulic conductivity in an ion-dependent manner (van Doorn *et al*., 2011; Santiago *et al*., 2013), and is relevant to understanding gas-liquid interactions in pit membranes.

For electrically charged walls that are not easily dehydrated, *h* is of the order of the Debye length (λ_D_), and has the approximate form:

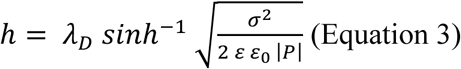

Here, sinh^-1^ is the inverse hyperbolic sine. Equation 3 is based on Hsu *et al*. (2003) and derived in the Supporting Information (SI) for a charged surface wall, but uncharged gas-liquid interface. Here, σ (expressed in C m^-2^, but practical units are frequently expressed in elementary charges per square nanometer, or e nm^-2^ = 0.1602 C m^-2^) is the surface charge density; ε (dimensionless; ε ≈ 80 at 20°C) is the relative permittivity or dielectric constant of water; ε_0_ (= 8.854 × 10⁻¹² C²·N⁻¹·m⁻²) is the vacuum permittivity; and |*P*| (expressed in Pa) is the magnitude of the negative pressure. Based on standard lipid biophysics, values of *σ* for a pit membrane coated with galactolipids and phospholipids (including phosphatidylcholine, phosphatidic acid, and phosphatidylinositol) can be estimated to vary from 0.1 to 0.5 e nm^-2^, which is considerably higher than estimations with cellulose/pectin surface chemistry (van Doorn *et al*., 2011; Santiago *et al*., 2013). Higher values of *h* are expected when the gas-liquid interface is also charged, in addition to the wall.

In essence, Equation 3 predicts how the equilibrium thickness of a water film adsorbed on a charged surface depends on the surface charge density and the negative pressure (Fig. S1). When the xylem sap pressure becomes more negative, it tends to draw water out of the film, making the equilibrium film thinner. This thinning is opposed by the repulsive electrostatic disjoining pressure generated by the charged surface and its diffuse ionic double layer, which stabilizes the remaining water film.

##### *R_eff_* = the effective pore radius

This radius is not a fixed structural parameter, but emerges from a hierarchical selection process in which gas invasion proceeds along the pathway that maximizes the minimum constriction radius (*R_min-max_*), after accounting for pressure-induced membrane deformation (*ΔR_mech_*). These dynamic and multiphase interactions make estimations of *R_eff_* challenging.

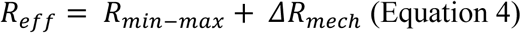

***R_min-max_*** is the largest of all minimal pore constrictions (see above), either in an individual pit membrane, or in all pit membranes of an interconduit pit field between two conduits, depending at what level *ΔP_bubble_* is estimated. This pore parameter can be estimated based on a stochastic model presented in Kaack et al. (2021), and is the original pore size before potential pit membrane deformation occurs. The value *R_min-max_* scales inversely and in a non-linear way with pit membrane thickness (Kaack *et al*., 2021).

The second component, ***ΔR_mech_*** is any change in pore constriction radius due to pressure-induced deformation. Cellulose microfibril aggregates in pit membranes are not interwoven (Pesacreta *et al*., 2005; Li *et al*., 2020), meaning that individual microfibril aggregates can move relative to each other under deformation. If a membrane bends or stretches under a pressure difference, changes in the pore geometry are likely, which may either widen or narrow local pore constrictions, i.e., resulting in positive or negative values, respectively. Deformation remains small at low pressure differences (Carmesin *et al*., 2023), but could increase nonlinearly with increasing pressure. *ΔR_mec_*_ℎ_ can be heterogeneous, and is likely largest at the centre of the pit membrane.

Estimating *ΔR_mech_* is challenging because it requires understanding how macroscopic displacement of pit membranes converts to microscopic changes in pore constrictions. A pore deformation factor (*χ*) could be introduced to link heterogeneous deformation of the multi-layered cellulose network (including local stretching, fibre rearrangement, pore-orientation, and anisotropic mechanical behaviour) with changes in the effective radius of the largest minimum pore constriction. The pore deformation factor *χ* is dimensionless (0 ≤ *χ* ≤ 1) and an emergent structural parameter that could be quantified in future work from high-resolution imaging or finite-element simulations.

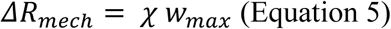

Importantly, *ΔR_mech_* relates to pit membrane displacement (w_max_), which can be estimated with the Kirchhoff-Love plate theory. Applying this approach, which quantifies the maximum central deflection of a pit membrane (Equation 6; Tixier *et al*., 2014), is reasonable because pit membranes can be assumed to be: (1) approximately circular and behave as isotropic elastic plates; (2) clamped at the pit border; and (3) relatively thin as compared to their diameter. The anisotropic, porous, and multi-layered nature of real pit membranes, however, suggest that applying the plate theory is a first-order approximation.

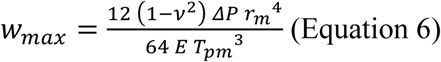

where *w_max_* (expressed in nm) is the maximal central deflection of the pit membrane; *ν* is the Poisson ratio (dimensionless), which can be estimated as 0.3 for biological, cellulose-rich, soft material (Tixier *et al*., 2014); *r_m_* is the radius of the pit membrane (µm); *E* is the Young’s modulus of a fresh, never-dried pit membrane, which was 57 MPa based on Carmesin *et al*. (2023); *ΔP* is the pressure difference across the pit membrane (MPa), and *T_pm_* is the pit membrane thickness.

Equations 5 and 6 suggest that a larger pit membrane with an increased radius undergoes dramatically more deflection and possibly higher values of *ΔR_mech_*, while an increase in pit membrane thickness strongly reduces deflection. For a pit membrane with a radius of 2 µm and pit membrane thickness of 300 nm, the maximum deflection becomes 530 nm under a pressure difference of 3 MPa. A slightly smaller pit membrane with a radius of 1.5 µm and a pit membrane thickness of 500 nm, however, will show a deflection of only 360 nm under the same pressure difference. When maximum deflection exceeds pit membrane thickness, nonlinear stretching and localized strain concentrations may become increasingly important, potentially promoting irreversible deformation and pore restructuring.

##### The broader relevance of the proposed embolism propagation concept

The dynamic concept of embolism propagation substantially changes our mechanistic understanding of embolism in plants. Most importantly, it revises several assumptions of the traditional air-seeding concept. The proposed two-step mechanism of embolism propagation also helps explain why plants have substantial resistance to embolism formation. Specifically, the combination of mesoporous pit membranes and polar, insoluble lipids lining xylem conduits indicate that bubble formation may occur without immediately triggering embolism. This concept introduces a dynamic, temporal component to embolism resistance, and helps explain how plants are able to transport xylem sap under negative pressure.

The dynamic nature of embolism propagation may affect how we measure embolism resistance of plants. As predicted by the traditional air-seeding hypothesis, the pressure gradient between embolized and water-filled conduits is the primary, although not exclusive determinant of embolism propagation. Gas diffusion rates, which are required for embolism propagation, are not only affected by the magnitude of this pressure gradient (Avila *et al*., 2023; Silva *et al*., 2024, 2025, 2026), but also by the duration of exposure to a given pressure difference, which adds a temporal component to vulnerability to embolism. Silva *et al*. (2024, 2026), for instance, showed that estimations of water potential values corresponding to 50% loss of hydraulic conductivity (i.e., P50 values) become on average 8.5% less negative for six angiosperm species in flow-centrifuge experiments that consider spin-time, while similar measurements of P50 without considering spin-time were more negative. The underlying mechanism is that gas diffusion through the xylem network to induce embolism formation can be relatively slow (Yang *et al*., 2023; Silva *et al*., 2025). Time also plays a role when the xylem sap from a recently embolised conduit provides a local buffering effect to further embolism spread (Hölttä *et al*., 2009; Vergeynst *et al*., 2015; Scheire *et al*., 2026). Several hours may be needed for a recently embolised conduit to build up atmospheric gas pressure, as gas movement within and between conduits relies mainly on axial and radial gas diffusion, with axial gas diffusion across interconduit pit membranes being ca. 100 times faster than radial diffusion across cell walls (Sorz and Hietz, 2006; Wang *et al*., 2015a, b; Yang *et al*., 2023). Moreover, temperature affects gas diffusion, and has a larger effect on gas solubility than pressure, with increasing temperature promoting gas exsolution from xylem sap (Mercury *et al*., 2003; Schenk *et al*., 2016). Yet, how temperature affects our estimations of embolism formation, especially with respect to gas solubility or dynamic surface tension of gas-liquid interfaces deserves more research (Silva *et al*., 2024).

Another example of how time may affect embolism resistance is that severe or recurring drought during the life span of a plant, may result in high nanobubble concentrations, potentially increasing the vulnerability to subsequent drought-induced embolism, a prediction that is consistent with the phenomenon of so-called “cavitation fatigue” (Hacke *et al*., 2001). It can be speculated that the accumulation of lipid-coated nanobubbles and/or polar lipids at the proximal end of xylem conduits over time could lead to a reduction of hydraulic conductivity, frequently observed in aging tree rings even in the absence of embolism (Wason *et al*., 2019; Fickle *et al*., 2025).

Moreover, mechanical stress, such as bending induced by wind, could cause xylem pressure changes that lead to embolism formation from existing nanobubbles, consistent with observations in *Citrus* trees (Michalczyk *et al*., 2025). The proposed separation of bubble propagation from embolism formation allows for old observations to be revisited and opens new avenues for research.

##### Open questions and future research priorities

New insights may provide opportunities to revisit interpretations of previous work, but also lead to new questions. Below, we discuss a few open questions that deserve further research.

Which conditions and processes determine the stability of surfactant-coated nanobubbles, and how exactly is embolism formation triggered from pre-existing bubbles? Because gas moves through pit membranes both in the form of lipid-coated nanobubbles and in dissolved form, with gas diffusion also taking place through cell walls (Silva *et al*., 2024; Sorz and Hietz, 2006), the equilibrium between these two gas phases critically affects the conditions required for embolism formation. Modelling of coated nanobubbles under changing negative pressure has shown that nanobubbles can be stable, but that there are low pressure conditions under which they will expand into embolism (Ingram *et al*., 2021, 2023, 2024). These models were run under yet another set of simplifying assumptions, and predictions of bubble behaviour depend on the nature of the lipids, the rate of change in temperature and pressure, and the concentration of dissolved gas. Moreover, dissolution of coated nanobubbles also appears possible under some conditions, and can be speculated to be associated with emission of acoustic signals.

It is unclear how the stability of nanobubbles is affected by the amount of dissolved gas in xylem sap, which can be over-saturated (Schenk *et al*., 2016). Based on Henry’s law, an equilibrium in the amount of gas dissolved in xylem sap would be expected and will be achieved over time as long as the system is not continuously driven away from equilibrium by temperature changes and incoming oversaturated sap (Schenk *et al*., 2016; Marion *et al*., 2026). Here, gas saturation refers to xylem sap containing dissolved gases at equilibrium solubility, whereas gas oversaturation refers to dissolved gas concentrations exceeding equilibrium solubility, creating the thermodynamic potential for bubble nucleation or growth. Gas oversaturation is likely due to relatively slow rates of (radial) gas diffusion (Sorz and Hietz, 2006), dynamic changes in various processes such as temperature, daily sap flow, respiration, and the uptake of soil water that is oversaturated with gas (Marion *et al*., 2026). Direct gas extraction of xylem sap shows daily patterns of gas concentrations that reflect dynamic phases of saturation to oversaturation (Pereira *et al.,* unpublished). Clearly, the dissolved gas saturation to oversaturation of xylem sap raises key questions on how plants can avoid constant failure of their hydraulic system by embolism.

The distributional, networking link behind embolism propagation provides interesting possibilities to model embolism spread in a similar way as an epidemic process using stochastic compartment models such as the susceptible-infected (SI) model, where the compartments correspond to sap-filled (“susceptible”) and embolised (“infected”) conduits (Roth-Nebelsick, 2019; Roth-Nebelsick and Konrad, 2024; Korhonen *et al*., 2026). Despite being a mesoscopic model that does not account for detailed pit anatomy, the SI model can capture embolism propagation accurately at the level of macroscopic xylem samples, with good agreement between the SI model, a physiological model, and experimental data (Korhonen *et al*., 2026). Among the family of compartment models, the SI model is conceptually close to the simplicity of air-seeding: embolism propagation between an embolised vessel and its sap-filled neighbour depends only on sap pressure, and any transmission event leads to immediate embolism of the sap-filled neighbour. On the other hand, a related compartment model, the susceptible-exposed-infected (SEI) model, is a conceptual match for the two-step phase of embolism propagation as proposed here. In this model, contact between embolised and sap-filled conduits leads first, with a certain probability, to exposure or formation of nanobubbles. The “exposed” conduits then only become embolised (“infected”) with a probability that corresponds to the nanobubble expansion. Simulations with compartment models emphasize the important effects of three-dimensional interconduit connectivity structure, as well as the connectivity structure between conduits, on embolism propagation (Loepfe *et al*., 2007; Mrad *et al*., 2018, 2021; Korhonen *et al*., 2026). This connectivity structure has been poorly studied despite the availability of methods such as microCT (Zimmermann and Tomlinson, 1966; Burggraaf, 1972; Wason *et al*., 2017, 2021). More attention is also needed to explore how the occurrence of tracheids affects vessel network properties and connectivity between conduits in angiosperms (Carlquist, 1984; Johnson *et al*., 2023).

Finally, there is more to learn about how bordered pits between conduits affect mechanical stability and protection of pit membranes, hydraulic efficiency, and safety from embolism. While all pits between conduits show a pronounced pit border, with considerable variation in bordered pit geometry and dimensions (Fig. 2; Sano *et al*., 2011; Kaack *et al*., 2019; Olson, 2023), it remains largely unclear why these interconduit pits are distinctly bordered. What are the functional consequences of the large variation in pit size, shape (from circular to oval and elongated), and arrangement (opposite, alternate, scalariform) on flow, potential pit membrane deformation, and bubble generation? The traditional idea that pit borders provide a mechanical support for aspirated pit membranes does not seem to apply to angiosperms based on the mechanical properties of fresh, never-dried pit membranes of *Clematis vitalba* (Carmesin *et al*., 2023), although this requires further research on more species. Moreover, species with vestured pits were found to show relatively thin pit membranes (Levionnois *et al*., 2021). For these thin, highly compliant pit membranes, the mechanical support provided by vestures acting as pillars remains a crucial hypothesis to prevent excessive displacement of the pit membrane (Fig. 2; Zweypfenning, 1978; Jansen *et al*., 2001, 2004; Choat *et al*., 2004; Medeiros *et al*., 2019; Levionnois *et al*., 2021).

## Conclusions

Recent findings on water transport in plants challenge key assumptions underlying the traditional air-seeding concept and lead us to conclude that this framework can no longer be interpreted as an accurate mechanism for embolism propagation between neighbouring conduits. Synthetizing key findings, we propose a new theoretical framework of embolism propagation to replace the traditional air-seeding concept by a two-step process of embolism propagation, which includes bubble generation and embolism formation as separate processes. Embolism propagation is governed by complex, multiphase interactions and dynamic processes that control how gas-liquid interfaces traverse interconduit pit membranes. A foam-like behaviour of pit membranes arises from the interplay between their dynamic porous structure, the surface activity of polar, insoluble lipids at gas-liquid interfaces, and snap-off processes within interconnected pore networks. Lipid-coated nanobubbles in bulk xylem sap can be stable, unless they become too large due to rapid changes in pressure, temperature, or nanobubble concentration. Embolism formation develops from these pre-existing bubbles, but not all bubbles lead directly to embolism formation. Consequently, the pressure threshold for gas-liquid interface movement is not static and cannot be described by a simple geometric pore constraint. Rather, it can be approximated by a modified, augmented Young-Laplace equation in which the effective pore radius is an emergent property, dynamically shaped by intrinsic pore architecture, and pressure-induced pit membrane deformation.

A two-step process of embolism propagation differs substantially from a one-step air-seeding concept, and has implications towards measurements of embolism resistance in plants, which may be affected by a time component. The new concept contributes to understanding why exactly plants can transport water under negative pressure without immediate and continuous formation of embolism in their conduits, which opens up interesting possibilities for transpiration-driven transport devices such as synthetic trees (e.g., Shi *et al*., 2020; Zweifel et al., submitted). Moreover, several new questions arise that warrant further investigation, particularly regarding the stability of nanobubbles under varying pressure, temperature, and gas saturation conditions; the ageing and mechanical properties of pit membranes (e.g., whether deformation induces porous-medium characteristics and alters pore-network properties); and the electrochemical properties of pit membranes. Recognising the new framework presented here will provide various opportunities for further work on how water transport in plants is affected by drought under global warming.

## Supplementary Information 1

### Glossary: list of terms used in the manuscript

**Air-seeding:** Embolism propagation between adjacent conduits as explained by Zimmermann’s (1983) hypothetical air-seeding mechanism.

**Apoplast, apoplastic:** All space in plant tissues that is not occupied by living cytoplasm. The apoplast includes cell walls, intercellular spaces, and the space inside dead xylem cells, such as fibres, tracheids, and vessels.

**Atmospheric pressure:** Pressure at the bottom of the atmosphere; approximately 0.1 MPa.

**Bordered pit:** A pit (i.e., an opening in a secondary cell wall) with a border and aperture that is considerably smaller than the pit membrane diameter.

**Bubble generation:** The formation of nanobubbles, which takes place in pit membranes.

**Bubble snap-off:** Bubble generation at pore constrictions within a pore pathway.

**Cavitation:** The nucleation of vapour bubbles in a liquid. Cavitation cannot occur under the pressure conditions in xylem.

**Conduit:** A dead xylem cell that transports sap. Conduits include tracheids and vessels.

**Constrictivity:** Indicator of pore constrictions.

**Debye-length:** The thickness of the electric double layer at a charged surface.

**Electric double layer:** The interfacial region formed by a charged surface and a surrounding layer of counterions in a liquid, resulting in a charge separation and an electrostatic potential gradient.

**Embolism formation:** The mechanistic process from expanding/coalescing nanobubbles in xylem sap to embolism of a conduit.

**Embolism propagation:** The process of spatial spread of embolism from one conduit to an adjacent one. Embolism propagation includes both bubble generation and embolism formation.

**Embolism:** The gas volume in a partially or completely gas-filled conduit.

**End wall:** The apical or basal wall of a conduit connecting to another conduit.

**Exsolution:** The process of gas coming out of solution into the gas phase.

**Fibres:** Lignified cells that do not transport sap but may store it. Can be dead or living. Fibres are characterised by simple to indistinctly bordered pits, and provide mechanical support.

**Imperforate tracheary elements:** Tracheary elements without a perforation plate. These include unicellular cells such as tracheids and fibres.

**Interconduit:** The cell wall or pit between two adjacent xylem conduits.

**Intervessel pit membrane area:** The total area of all intervessel pit membranes on a vessel wall.

**Mesoporous:** A porous medium with pores between 5 and 50 nm in diameter.

**Nanobubble:** Here defined as a surfactant-coated gas bubble between 20 and 300 nm in radius. Nanobubbles of this size without surfactant coats are not stable in water.

**Negative pressure:** Absolute pressure below 0 MPa, only possible in liquids and solids, used here as a synonym for tension.

**Permeability:** The intrinsic ability of a porous medium to transmit fluid through its interconnected pore space. In an interconduit pit membrane, permeability is a structural property that describes how readily xylem sap can pass through the pit membrane’s pore network between adjacent conduits.

**Pit aperture:** The opening in the pit border, with the inner pit aperture facing the lumen of the conduit, and the outer pit aperture facing the pit chamber.

**Pit border:** The overhanging secondary cell wall, which partially encloses the pit chamber.

**Pit canal:** The canal between the inner pit aperture and the pit chamber; it is as long as the border is thick or sometimes longer if the canal is not straight.

**Pit chamber:** A semi-enclosed volume surrounded by the roof of the overhanging pit border and the pit membrane.

**Pit membrane area:** The pit membrane area of an individual pit.

**Pit membrane:** The primary cell wall in a pit. In case of interconduit, bordered pits, the pit membrane becomes chemically modified during programme cell death of the conduits.

**Pith:** The central tissue of a plant stem or root.

**Pore constriction:** A locally narrowed space within a three-dimensional pore pathway. The term pore throat is a synonym of pore constriction.

**Pore network:** The entire connected ensemble of pores and voids in a porous medium, representing all structurally possible connectivity routes under hydrated conditions. In an interconduit pit membrane, the pore network is a three-dimensional assembly of nanopores and constrictions, where many overlapping pathways can potentially connect adjacent pit borders.

**Pore pathway:** One specific, continuous route through a pit membrane, composed of interconnected pore spaces and pore constrictions. It represents a transient, hydraulically continuous percolation pathway that allows xylem sap or gas to transverse the pit membrane under specific pressure and interfacial conditions.

**Porosity:** The pore volume fraction of a porous medium. It quantifies how much pore space exists, and not how large the pores are. Porosity is frequently misused as describing the pore size of a porous medium.

**Surface tension, bulk:** The surface tension at a gas-liquid interface in the absence of insoluble surfactants, depending on the nature of the liquid and dissolved surfactants.

**Surface tension, dynamic:** The surface tension at a gas-liquid interface, depending on the local concentration of insoluble surfactants, such as polar lipids, per surface area. Dynamic surface tension varies in both space and time.

**Surface tension, equilibrium:** The dynamic surface tension at a gas-liquid interface in the absence of forces that stretch or compress insoluble surfactants.

**Tension:** See negative pressure.

**Tortuosity, geodesic:** The ratio of the mean shortest connected path length to the length of the porous medium; here this length is the pit membrane thickness.

**Tracheid:** Unicellular conduit with bordered pits and without perforation plate.

**Vessel:** Multicellular conduit with bordered pits, characterised by perforation plates between vessel elements.

**Vestures, vestured:** Outgrowths from the pit border into the pit chamber. These can be limited to small, unbranched outgrowths near the pit aperture, or distinctly branched vestures could fill up the entire pit border.

**Wettability:** the tendency of a liquid to spread on or adhere to a solid surface (e.g., conduit wall or pit membrane) in the presence of a gas phase. It is commonly quantified by the contact angle at a solid-liquid-gas interface.

## Supplementary Information 2

### S1 Electrostatically stabilised water film on a charged surface

To estimate the electrostatic contribution to the critical pressure for bubble generation, we calculate the equilibrium thickness of a water film that remains adsorbed on a charged surface after the passage of an air front (Figure S1A). This film is stabilized by an electrostatic disjoining pressure *Π_el_*(ℎ), which provides the additional pressure term used in the augmented Young-Laplace equation in the main text.

We consider a planar substrate carrying a uniform fixed surface charge density *σ* and covered by a water film of equilibrium thickness *h*, whose outer boundary is the air-water interface.

To determine *h*, we first consider the classical electrostatic problem of two parallel charged surfaces carrying uniform surface charge densities *σ1* and *σ2*. Assuming that the electrostatic potential remains sufficiently small, it satisfies the linearized Poisson-Boltzmann (Debye-Hückel) equation,

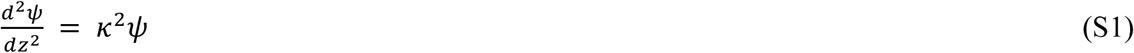

where *κ⁻¹* = *λ_D_* is the Debye screening length.

The electrostatic potential satisfies the following boundary conditions (Holm et al., 2000):

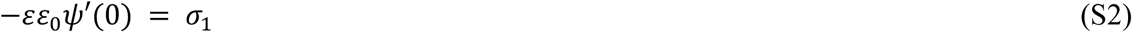

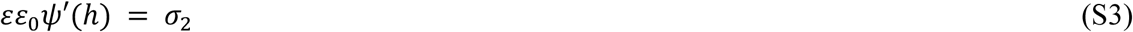

where *ε* ≈ 80 is the relative permittivity of water.

The corresponding electrostatic disjoining pressure is given by Hsu et al. (2003):

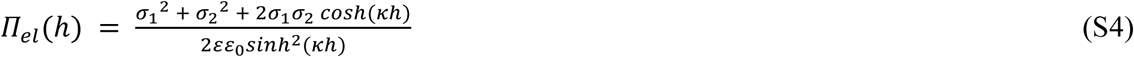

In the present system, one boundary corresponds to the charged constriction wall, whereas the opposite boundary is the air-water interface, which is assumed here not to be charged. Setting *σ1* = *σ* and *σ2* = 0 therefore yields

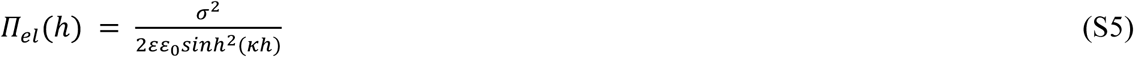

In mechanical equilibrium, the electrostatic disjoining pressure balances the applied hydrostatic pressure (*P* < 0),

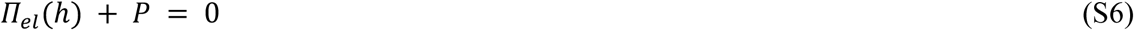

Non-electrostatic contributions to the electrostatic disjoining pressure, including van der Waals and hydration forces, as well as the air pressure, are neglected.

Solving Eq. (S6) yields the equilibrium water-film thickness,

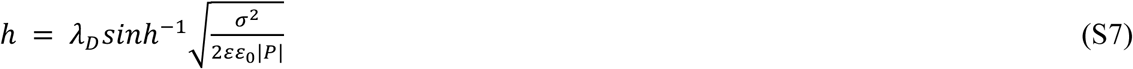

where sinh⁻¹ denotes the inverse hyperbolic sine function. Equation (S7) predicts that the film thickness increases with increasing surface charge density and decreases as the magnitude of the negative pressure increases. Figure S1B shows the equilibrium thickness, *h*/*λ_D_*, as a function of the surface charge density.

**Figure S1:**
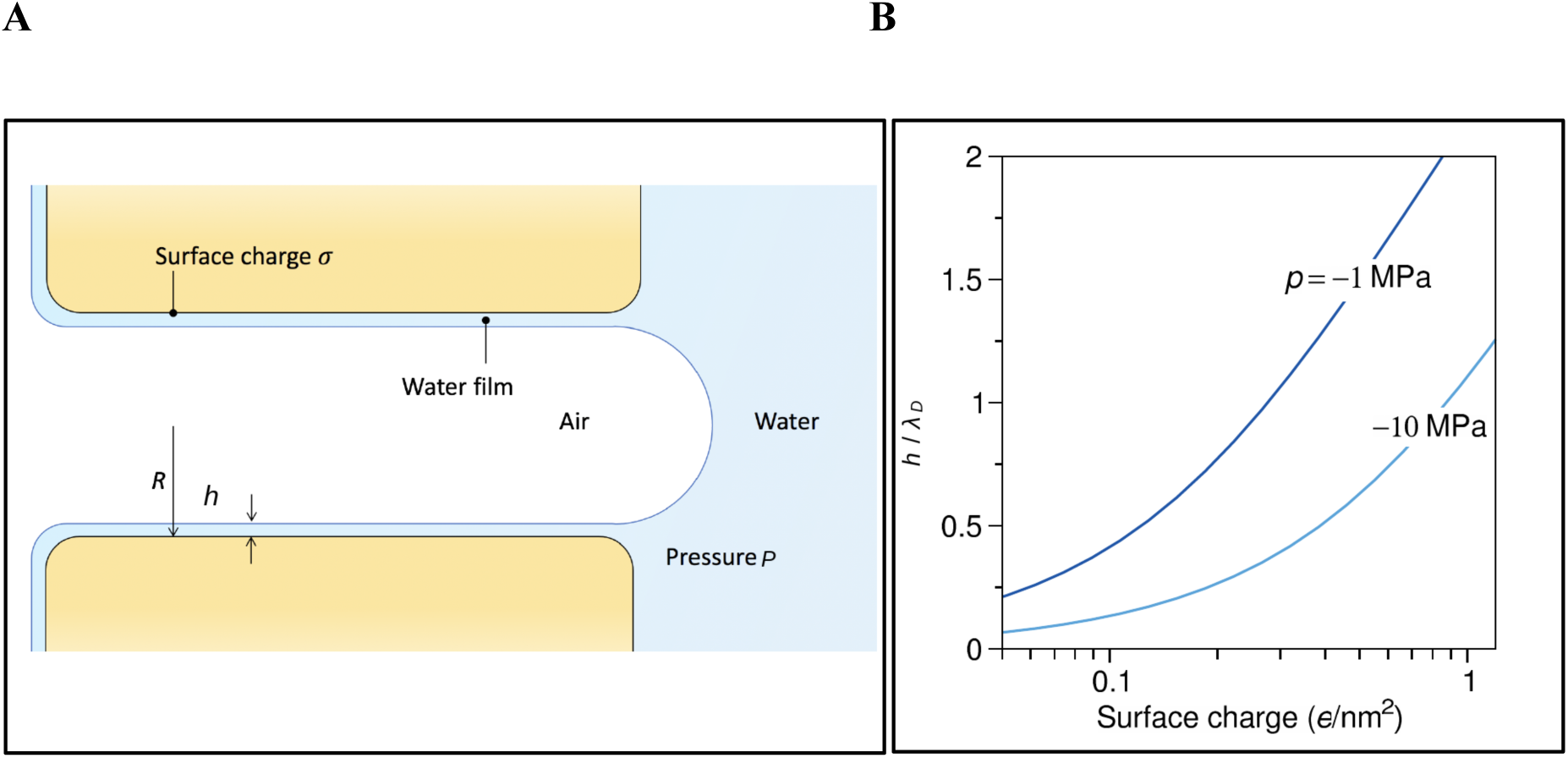
**A.** Schematic illustration of a pore constriction with a uniformly charged surface (surface charge density *σ*; which is quantified as elementary charges per square nanometer or e nm^-2^ = 0.1602 C m^-2^), which stabilizes an adsorbed water film of thickness *h*. The film is in contact with bulk water under a negative pressure *P*. *R* is the radius of the pore constriction. **B.** Dimensionless equilibrium film thickness, *h*/_λD_, as a function of surface charge density for two values of hydrostatic pressure *P*, calculated from equation S7.

## Acknowledgements

We thank the Electron microscopy section of Ulm University for technical support over the last 17 years. Valuable comments from Jordi Martínez Vilalta on an earlier version of this manuscript were highly appreciated.

